# ACE2 interaction networks in COVID-19: a physiological framework for prediction of outcome in patients with cardiovascular risk factors

**DOI:** 10.1101/2020.05.13.094714

**Authors:** Zofia Wicik, Ceren Eyileten, Daniel Jakubik, Sérgio N Simões, David C Martins, Rodrigo Pavão, Jolanta M. Siller-Matula, Marek Postula

## Abstract

**Background:** Severe acute respiratory syndrome coronavirus 2 (SARS-CoV-2) infection (coronavirus disease 2019; COVID-19) is associated with adverse outcomes in patients with cardiovascular disease (CVD). The aim of the study was to characterize the interaction between SARS-CoV-2 and Angiotensin-Converting Enzyme 2 (ACE2) functional networks with a focus on CVD.;

**Methods:** Using the network medicine approach and publicly available datasets, we investigated ACE2 tissue expression and described ACE2 interaction networks which could be affected by SARS-CoV-2 infection in the heart, lungs and nervous system. We compared them with changes in ACE-2 networks following SARS-CoV-2 infection by analyzing public data of stem cell-derived cardiomyocytes (hiPSC-CMs). This analysis was performed using the NERI algorithm, which integrates protein-protein interaction with co-expression networks. We also performed miRNA-target predictions to identify which ones regulate ACE2-related networks and could play a role in the COVID19 outcome. Finally, we performed enrichment analysis for identifying the main COVID-19 risk groups.

**Results:** We found similar ACE2 expression confidence levels in respiratory and cardiovascular systems, supporting that heart tissue is a potential target of SARS-CoV-2. Analysis of ACE2 interaction networks in infected hiPSC-CMs identified multiple hub genes with corrupted signalling which can be responsible for cardiovascular symptoms. The most affected genes were EGFR, FN1, TP53, HSP90AA1, and APP, while the most affected interactions were associated with MAST2 and CALM1. Enrichment analysis revealed multiple diseases associated with the interaction networks of ACE2, especially cancerous diseases, obesity, hypertensive disease, Alzheimer’s disease, non-insulin-dependent diabetes mellitus, and congestive heart failure. Among affected ACE2-network components connected with SARS-Cov-2 interactome, we identified AGT, CAT, DPP4, CCL2, TFRC and CAV1, associated with cardiovascular risk factors. We described for the first time miRNAs which were common regulators of ACE2 networks and virus-related proteins in all analyzed datasets. The top miRNAs were miR-27a-3p, miR-26b-5p, miR-10b-5p, miR-302c-5p, hsa-miR-587, hsa-miR-1305, hsa-miR-200b-3p, hsa-miR-124-3p, and hsa-miR-16-5p.;

**Conclusion:** Our study provides a complete mechanistic framework for investigating the ACE2 network which was validated by expression data. This framework predicted risk groups, including the established ones, thus providing reliable novel information regarding the complexity of signalling pathways affected by SARS-CoV-2. It also identified miR which could be used in personalized diagnosis in COVID-19.

## 1. Introduction

At the end of 2019 in Wuhan (China), a novel coronavirus named severe acute respiratory syndrome coronavirus 2 (SARS-CoV-2) has been discovered [1]. The clinical manifestations of SARS-CoV-2 infection, named coronavirus disease 2019 (COVID-19), varies in severity from asymptomatic infection to acute viral pneumonia with fatal outcome. Nearly half of patients who were at risk of acute course of the disease suffered from comorbidities including hypertension, diabetes mellitus (DM) and coronary heart disease [2,3]. Importantly, COVID-19 is associated with increased risk for mortality and adverse cardiovascular events among patients with underlying cardiovascular diseases (CVD) [4]. A similar association between the virus and CVD was observed during previous coronavirus outbreaks caused as Middle-East respiratory syndrome coronavirus (MERS) or severe acute respiratory syndrome coronavirus (SARS-CoV) [5,6]. Therefore, these data suggest a common factor that is associated with the pathogenesis of COVID-19 and CVD. The etiology of cardiac injury in COVID-19, however, still remains unclear. It is hypothesized that cardiac injury may be ischemia mediated, and the profound inflammatory and hemodynamic impacts are seen in COVID-19 can cause atherosclerotic plaque rupture or oxygen supply-demand mismatch resulting in ischemia [7]. Most probably, the link between cardiovascular complications and infection may be related to angiotensin-converting enzyme 2 (ACE2), which was found to act as a functional receptor for SARS-CoV-2 [8].

ACE2 is a multi-action cell membrane enzyme that is widely expressed in the lungs, heart tissue, intestine, kidneys, central nervous system, testis, and liver [9]. During the 20 years from its discovery, the investigations targeting the complex role of this enzyme established ACE2 as an important regulator in hypertension, heart failure (HF), myocardial infarction (MI), DM, and lung diseases [10,11]. The viral entry to cells is determined by the interaction between the SARS-CoV-2 spike (S) protein and N-terminal segment of ACE2 protein, with a subsequent decrease in ACE2 surface expression, which may be enhanced by cofactor transmembrane protease serine 2 (TMPRSS2) [12]. Publications of independent research groups showed that cardiomyocytes can be infected by SARS-CoV-2. The virus can enter hiPSC-CMs via ACE2, and the viral replication and cytopathic effects induce hiPSC-CM apoptosis and cessation of beating after 72 h of infection while inhibiting metabolic pathways and suppressing ACE2 expression during this initial infection stage [13]. Moreover, SARS-CoV-2 undergoes a full replication circle and induces a cytotoxic response in cardiomyocytes, by inducing pathways related to viral response and interferon signalling, apoptosis and reactive oxygen stress [14]. Consistently, mRNA and protein ACE2 are expressed in hiPS-CM, whereas TMPRSS2 was detected only at very low levels by RNA sequencing [14]. Finally, SARS-CoV-2 was invariably detected in cardiomyocytes of COVID-19 patients without clinical signs of cardiac involvement, with degrees of injury ranging from the absence of cell death and subcellular alterations hallmarks to intracellular oedema and sarcomere ruptures [15]. These findings support that heart tissue can be infected with SARS-Cov-2.

As a consequence of SARS-CoV-2 infection, downregulated ACE2 pathways may lead to myocardial injury, fibrosis, and inflammation which may be responsible for adverse cardiac outcomes [16]. In line with these findings, several reports linked SARS-CoV-2 infection with myocardial damage and HF, accompanied by acute respiratory distress syndrome (ARDS), acute kidney injury, arrhythmias, and coagulopathy. The incidence of myocardial injury ranged from 7 to 28% depending on the severity of COVID-19, accompanied by increased levels of cardiac troponins, creatinine kinase–myocardial band, myohemoglobin, and N-terminal pro-B-type natriuretic peptide (NT-proBNP) [17–19].

In the current work, we characterized the ACE2 interaction network in the context of myocardial injury. Our quantitative *in silico* analysis pointed out: (1) the potential tissues and organs which can be infected by SARS-CoV-2; (2) the top ACE2 interactors associated with the virus-related processes with altered co-expression networks in hiPSC-CMs after SARS-CoV-2 infection, which likely play role in the development of CVD; (3) signalling pathways associated with alteration of ACE2 networks; (4) prediction of risk groups in COVID-19; (5) connections between ACE2 and SARS-Cov2 interactomes, as well as ACE2 co-expression networks in hiPSC-CMs; (6) the most promising microRNAs (miRNAs, miR) regulating ACE2 networks for potential diagnostic and prognostic applications.

Our comprehensive analysis investigating ACE2 receptor-related interaction networks, their connection with SARS-CoV-2 interactome, enriched signalling pathways, miRNAs, associated diseases, provide precise targets for developing predictive tools, with the potential for reducing health, personal and economic consequences of the pandemic.

## 2. Materials and Methods

### 2.1. Data collection

ACE2-associated genes used for constructing interaction networks were extracted from KEGG database (23 genes from renin-angiotensin system-RAS pathway) [20]; stringApp (top 40 ACE2 interactors) [21], Archs4 database https://amp.pharm.mssm.edu/archs4 (top 20 genes with correlated expression), Genecards database https://www.genecards.org (5 interactors and 4 sister terms), literature search [22–24]. In total, we collected 69 genes, which were used further for miRNA prediction analysis and constructing interaction networks. In all steps of data integration and bioinformatic analyses, we used our R package wizbionet [25]. Tissue Expression analysis

The tissue expression of ACE2 and TMPRSS2 was evaluated based on a dataset downloaded from the Tissues 2.0 database and the GTEX database [26]. Tissues 2.0 database integrates: transcriptomics datasets from multiple sources and proteomics datasets from humans and other organisms, quantifying gene expression confidence scores across tissues. All tissues were sorted by the decreasing expression confidence of ACE2 and TMPRSS2. Additionally, mean expression confidence for the ACE2 network was calculated for each tissue from the database. Gene expression confidence scores were also mapped on the visualization of the interaction networks.

### 2.2. Interaction network analysis

#### 2.2.1. ACE2 interactome

To analyze connections between ACE2 and other genes, we constructed a PPI network in Cytoscape 3.7.2, using human interactome data from StringApp 1.5.1 (Search Tool for the Retrieval of Interacting Genes/Proteins) database, including known and predicted protein-protein interactions [21,27]. Interaction networks were composed of a set of genes (nodes) connected by edges that represent functional relationships among these genes. As suggested by the StringApp, we took into account connections with edge interaction confidence cut-off >0.4 (medium confidence), with 1 being the highest possible confidence and 0 lowest. We compared the complete tissue-specific ACE2 network, across the heart, lungs, and nervous system, as well as the virus-infection-related proteins network. The criteria of selection tissue-specific networks were gene expression confidence score >2.

#### 2.2.2. NERI Method

NERI (NEtwork by Relative Importance) [28] is a method that computes the relative importance of genes related to the seeds. The method is based on the Network Medicine Hypotheses: Locality, Disease Module, and Network Parsimony. It integrates PPI network with expression data (from the two conditions, *e*.*g*. control and disease), and uses a previously chosen seed genes list. NERI computes two relative importance scores for each gene, one score for each expression condition (control and disease). This is done by selecting the best shortest paths (based on the Parsimony Hypothesis) from seeds to their neighborhood (based on Locality and Disease Module Hypotheses) and taking into account the expression condition. The adopted criterion to evaluate a path is the modified Kendall’s concordance (a way to measure a group correlation) of expression of genes in the given path -- the more concordant a path is, the better. Then, the relative importance score of a given gene is the sum of all concordances of the selected paths to which the gene belongs, weighted by the proximity to the seeds. This procedure is executed for two conditions independently, generating two networks (e.g. control and disease networks). In the end, the NERI method performs the differential network analyses, which outputs two ranked lists of genes (*X, Δ’*): one based on the sum of the relative importance scores, and another based on the normalized difference between the relative importance. The first one (X) prioritizes genes with party hub features, possessing high topological centrality and, at the same time, high co-expression relative to the seed genes. The second one (Δ’) prioritized the most altered genes between two conditions as described before [28]. In the present study, we used the NERI algorithm to analyze raw expression signals obtained from the GSE150392 dataset. Besides, we used the BIOGRID dataset to construct the PPI network and, as seeds genes, the 69 ACE2-related interactors collected as described in the method section. Differential expression analysis for visualization was performed using the Mann Whitney test with p FDR corrected <0.05.

### 2.3. Extraction of disease-relevant ontological terms

To improve the interpretation of the gene functions, we mined Gene Ontology (GO) database using the biomaRt R package for extracting GO terms and further genes associated with the following processes: “inflammation” (22 GO terms; 648 genes), “coagulation” (18 GO terms; 223 genes) [29], “angiogenesis” (24 GO terms; 535 genes), “cardiac muscle functions” (176 GO terms; 524 genes), “muscle hypertrophy” (16 GO terms; 85 genes), and “fibrosis” (23 GO terms; 263). A similar methodology was used to extract genes potentially related to “viral infection” (120 GO terms, 1047 genes). Disease relevant gene lists were extracted from the http://t2diacod.igib.res.in/ database [30]. From this database, we used atherosclerosis, nephropathy, CVD, and neuropathy datasets. Diabetes-related genes were extracted from the StringApp disease database, for the term “diabetes type-2”. GO term lists used for gene extraction are shown in [Supplementary Table 1](https://ln.sync.com/dl/1d778f4a0/93k5i3kk-fekza6km-jize2twa-k5cu3f7w).

### 2.4. Enrichment analysis

Enrichment analysis is a computational method for increasing the likelihood to identify the most significant biological processes related to the study [31]. Enrichment analysis of the diseases and networks was done with the EnrichR database [32], using Fisher’s exact with Benjamini and Hochberg correction, while the reference was precomputed background for each term in each gene set library. Signalling pathways were analyzed using BioPlanet2019 and Human KEGG 2019 datasets. Diseases were analyzed using DisGenet Dataset and are Diseases AutoRIF Gene Lists datasets. In all statistical analyses, the significance cutoff was set to corrected p-value ≤0.05.

### 2.5 miRNA predictions

In order to identify miRNAs regulating ACE2 related genes, we used R and multimiR package with default settings, similar to previous publications from our group [29,33–35]. Interaction networks between ACE2-related genes and miRNAs were constructed in R and exported to Cytoscape 3.7.2. Next, the interaction networks were merged with the predicted PPI network for ACE2 constructed using StringApp. Both networks were merged using official gene symbols and ENSG numbers.

## 3. Results

### 3.1. ACE2 tissue-specific expression

To evaluate the potential susceptibility of the heart for SARS-CoV-2 infection, we ranked all 6668 human tissues from the Tissue 2.0 database, based on provided by the database ACE2 expression confidence (scale 0-5). This analysis revealed that lungs and respiratory systems are in the 14th and 15th place in terms of ACE2 expression confidence, after heart and cardiovascular systems (12th and 13th place, respectively), and before the nervous system (16th place; Figure 1 A). TMPRSS2 gene expression confidence was lower in the lungs and heart than in the nervous system. The mean expression of 69 genes from the network was highest in the nervous system in comparison to heart and respiratory system tissues. Exceptionally high scores for ACE2 expression were assigned to urogenital and reproductive tissues. As an additional validation, we used the expression dataset from the GTEX database, which confirmed our findings using the TISSUES2.0 database (Figure 1 B). According to the GTEX database testis showed the highest expression of ACE2, while the prostate showed high expression of TMPRSS2 Heart-related tissues located respectively in 5, 7, and 15th place. In turn, the gastrointestinal tract in GTEX appears in 2ndplace while in the Tissues 2.0 database it was not present among top hits. In both datasets, female reproductive glands (Tissues2.0) and mammary glands (GTEX) showed a high expression of the ACE2 receptor.

**Figure 1.**
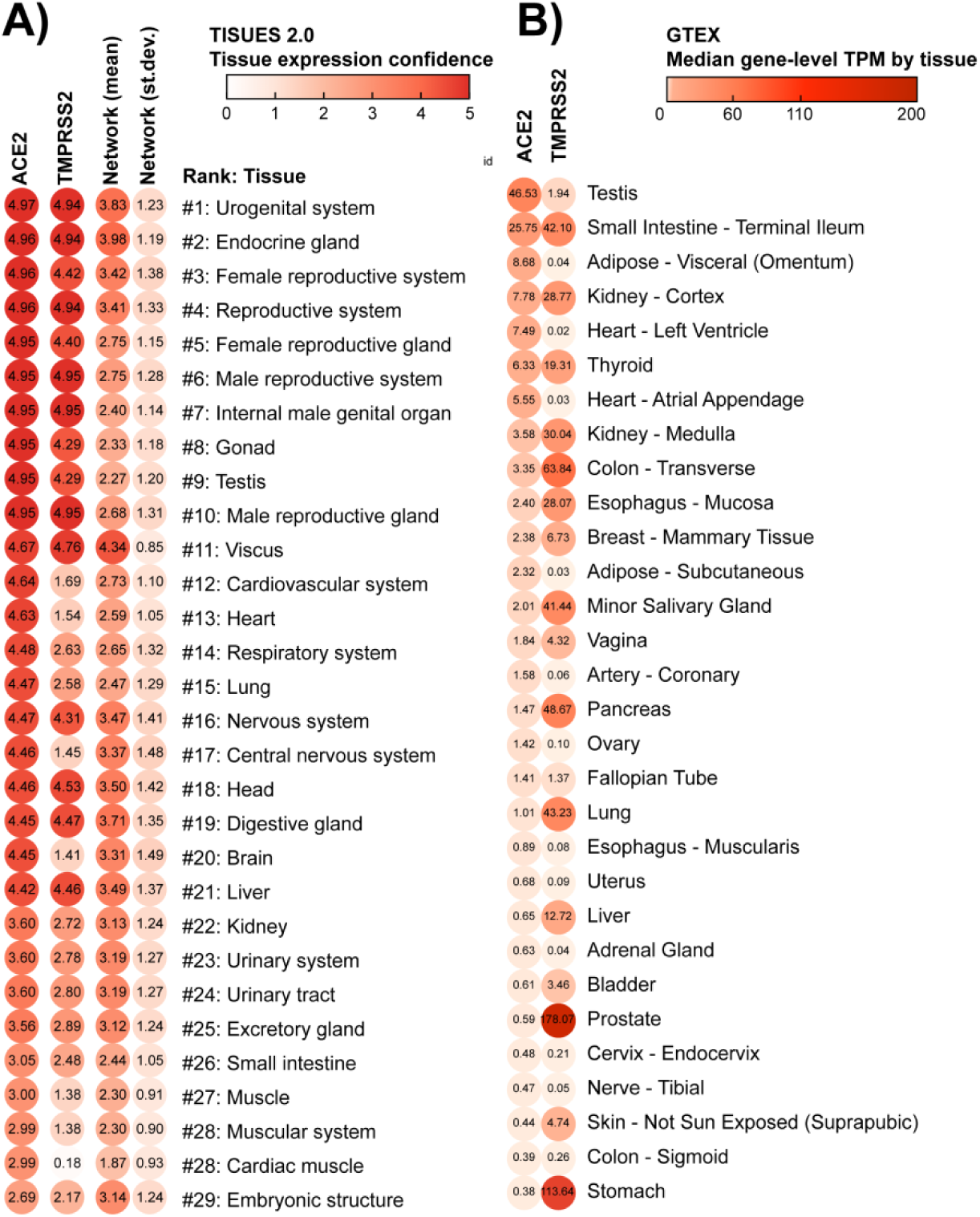
Tissues sorted by the potential of being infected by SARS-CoV-2. These lists of tissues were generated according to the concentration of membrane receptors ACE2 and TMPRSS2, obtained from (A) TISSUES 2.0 database expression confidence values, and (B) GTEX database Transcripts Per Million (TPM). The virus starts the cell infection by binding to ACE2, a major hub in multiple physiological processes: this binding can block ACE2 network activity. However, the virus will enter the host cell when TMPRSS2 cleavages ACE2. The first column depicts the average gene and protein expression confidence for the ACE2 receptor; the second column depicts the average expression confidence of TMPRSS2. The mean and standard deviation of expression confidence across 69 genes/proteins of the ACE2 network are presented in third and fourth columns, respectively. Notice that the lungs and respiratory system are ranked as #14-15 in the TISSUES 2.0 list, while the heart and cardiovascular system #12-13. Nervous and reproductive systems are ranked as, #16-17 and #1-10, respectively.

### 3.2. Construction of the complete ACE2 network

By using available literature and interactome data, we made an attempt to construct the ACE2 network as complete as possible based on the interactions with other genes and proteins. We developed the network based on three assumptions from the network medicine field: (i) disease module hypothesis: gene-products associated with the same disease phenotype tend to form a cluster in the PPI network; (ii) network parsimony: shortest paths between known disease genes often coincide with disease pathways; (iii) local hypothesis: gene products associated with similar diseases are likely to strongly interact with each other [36]. Besides protein-protein interactions, we also included genes that showed correlated expression with ACE2, taking into account that these genes could not yet have a strong representation in PPI databases. The aim of this synthesis was to gather available knowledge regarding ACE2 interactome which could be useful for interpreting new findings in the context of the disease and could be further narrowed down by using their own expression or proteomic data. In our work, we used expression data from Tissues 2.0 for subsetting tissue-specific sub-networks and Gene Ontology to identify virus specific-proteins. This complete ACE2 network also provided a starting point for our new analysis added to the manuscript of SARS-CoV-2 infected hiPSC-CMs, where those 68 genes served as a seed nodes for the NERI algorithm which integrated co-expression networks with PPI networks.

From 68 genes included to complete the ACE2 network, only two didn’t show any interactions according to the String database which we used for the visualization of the network (Figure 3 A). This figure depicts the complexity of this disease, presenting the possible primary alterations following SARS-CoV-2 infection, on proteins level, and the secondary alterations, on gene expression level (including initial ACE2 downregulation and later ACE2 overexpression). The genes from the ACE2 network were sorted using circular sorting by the number of connections with other genes to simplify the visualization. The workflow of the bioinformatic analyses performed in this study is shown in Figure 2.

**Figure 2.**
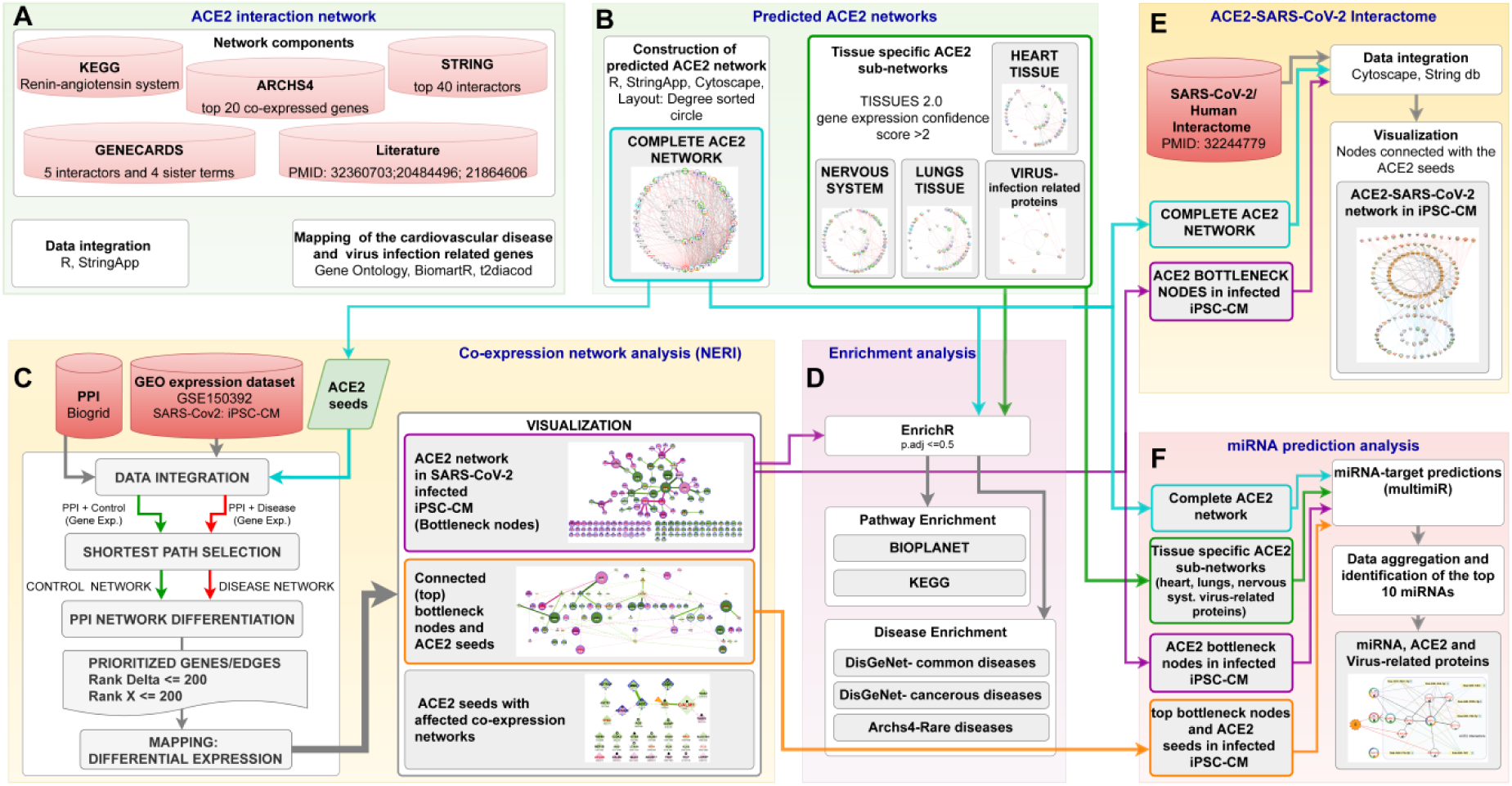
The workflow of bioinformatic analyses. (A) data collection to construct complete ACE2 network; (B) generation of the complete ACE2 network as well as tissue-specific sub-networks; (C) ACE2-related co-expression network analysis of pluripotent stem cell-derived cardiomyocytes (hiPSC-CMs) 72h post-infection with SARS-CoV2 using NERI algorithm; (D) Enrichment analysis of signalling pathways and diseases related to alterations in ACE2 networks; (E) integration of complete ACE2 network with NERI; and (F) miRNA prediction analysis in ACE2 related networks.

**Figure 3.**
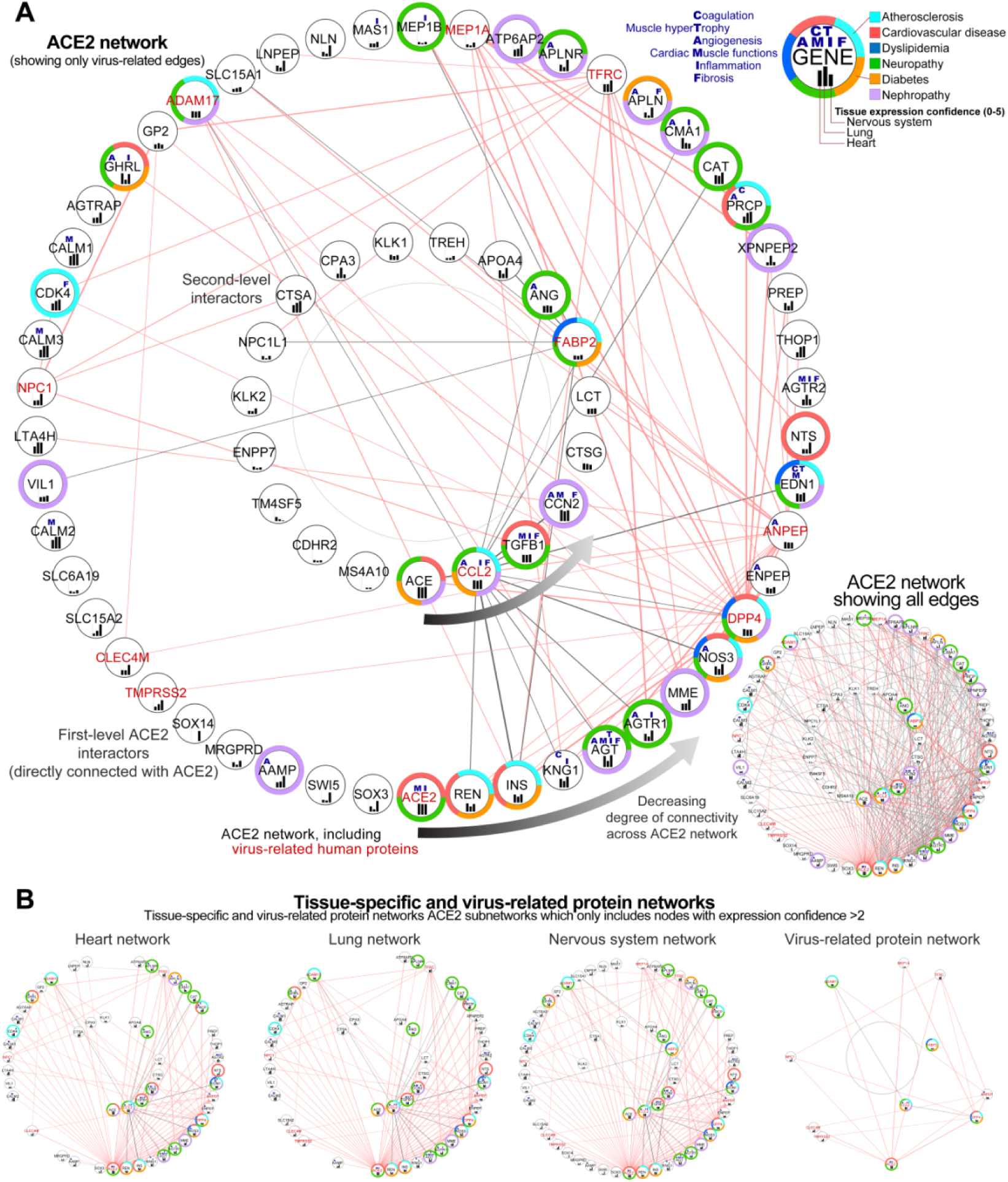
Predicted ACE2 interaction network. (A) Complete ACE2 network visualized as two circles sorted by the number of connections (degree) with other nodes. The external circle depicts the first level ACE2 interactors; the internal circle depicts the second level of interactors, with genes that do not connect directly with ACE2. For clarity, on the main figure, we showed only the edges associated with virus-related proteins (gene id in red). Edges associated with virus-related proteins are shown in red for first-level (direct) and in grey for the second-level (indirect) ACE2 interactors. Inset in the top right depicts the additional information for each gene/protein, as associated processes (blue letters), associated diseases (color-label ring), and expression confidence across key tissues (black bars). Inset in the bottom right depicts the same network including all edges. The genes from the ACE2 network were sorted using a circular layout organizing the genes by the number of connections with other genes which enabled us to hide the edges to simplify visualization. Thus, genes present on the bottom toward the right of the circular network showed the highest connectivity within the network. Notice that the closest ACE2 interactors are ACE, REN and INS, which play a central role in the pathophysiology of a number of cardiovascular disorders. The following interactor is KNG1, essential for blood coagulation and assembly of the kallikrein-kinin system and AGT influencing the renin-angiotensin system (RAS) function. In the network are present 11 virus-infection-related proteins (red labels) forming a dense connection with ACE2 and its top interactors which can affect its functionality. (B) Subsets of ACE2 network containing only highly expressed proteins in the heart, lung, and nervous system; analogous network for virus-related proteins (right). From the genes which didn’t have direct interactions with ACE2, the gene ACE showed the highest connectivity.

#### 3.2.1. Identification of genes showing the highest connectivity within complete ACE2 network

Analysis of the complete interaction network between ACE2 and associated genes showed, as expected, the highest number of interactions between ACE2 and other genes (49 interactions), followed by ACE, which was not directly connected with ACE2 (33 interactions), renin (REN, 32 interactions), insulin (INS, 31 interactions), kininogen 1 (KNG1, 30 interactions) and angiotensinogen (AGT, 28 interactions) (Figure 3 A).

##### Identification of the virus infection-related proteins within complete ACE2 network

Analysis of the ACE2 interaction network revealed 11 genes associated with virus infection-related ontological terms (ACE2, DPP4, ANPEP, CCL2, TFRC, MEP1A, ADAM17, FABP2, NPC1, CLEC4M, TMPRSS2) which could be especially affected in SARS-CoV-2 infection, leading to disturbance of the network. All these genes were connected directly with ACE2 according to the String database, except for gene FABP2 (6 interactions with other genes) and CCL2 (15 interactions). From this group, the highest degree of connectivity with other genes from the network was found for DPP4 (22 interactions), ANPEP (19 interactions), and CCL2 indirectly connected with ACE2 (15 interactions, as presented before). Two genes CDHR2 and MS4A10 didn’t have any known connections with other genes according to the String database, for edge confidence score cut-off >0.4.

#### 3.2.2. Sub-setting of the tissues-specific ACE-2 related networks

We also made subsets of complete ACE2networkto show interactions for heart tissue, lungs, nervous system as well as virus-infection related proteins. Tissue-specific networks were selected based on their tissue expression confidence in analyzed tissues. These enabled us to evaluate similarities between selected tissues to predict the impact of ACE2 alterations in the heart tissue. Lung tissue was selected based on how affected it is by the SARS-CoV-2 infection and served as the control for our analysis. Nervous tissue was selected due to multiple neurological symptoms recently reported as associated with COVID19 disease [37].

We set this score as 2, which is relatively high taking into account scores for the genes from this database. For all tissues, the median expression confidence score was 0.914, for the heart was 0.989, lung 1.319, nervous system 1.556. We found 48 genes from 68 overlapped between tissue-specific networks, 8 of them were virus-infection related proteins (Fig 3 B). This ACE2-interactome also provided a starting point for our analysis of ACE2 coexpression networks in cardiomyocytes performed in this study, where those 68 genes served as seed nodes for the NERI algorithm which integrated co-expression networks with PPI networks.

Descriptions of the genes from the complete ACE2 network and link to its interactive version are available in [Supplementary Table 2](https://ln2.sync.com/dl/80b495830/4whdyy4r-phxq25b2-eqemr3ei-ap37by6u/view/doc/8923100690005).

### 3.3. Analysis of changes in ACE2 co-expression network in infected cardiomyocytes

To evaluate how alteration in the ACE2 network can affect cardiomyocytes we re-analyzed the GSE150392 GEO dataset for pluripotent stem cell-derived cardiomyocytes (hiPSC-CMs) 72h post-infection with SARS-CoV2. We utilized algorithm NERI [28],[38] that integrates the PPI interactome data with co-expression networks to take a closer look at this part of the cardiomyocyte co-expression network. We used 68 genes from the predicted ACE2 network as seed genes, to focus on this region of the transcriptome. The goal of this analysis was to identify the genes and interactions between them that could be affected by the alterations in the ACE2 protein network caused by the virus and impact the functionality of cardiomyocytes. By assuming some network medicine hypothesis, the method explored the neighborhood of a gene set by locating paths possessing more coexpressed genes with seeds - this is independently performed for two conditions (control and disease). This approach enabled us to identify a cluster of hub genes also called “bottleneck regulators”, with corroborating signals across transcript and protein expression data leading to pathological changes in cardiomyocytes even when they show little or no changes in expression. In order to identify those genes, we selected nodes and edges in which Rank scores generated by NERI ranged from 1 to 200 in terms of the difference in co-expression between control and disease (Rank Delta) and changes in connectivity (Rank X).

#### 3.3.1 Identification of the ACE-2 related bottleneck genes related to corrupted co-expression networks

We identified 139 bottleneck genes, four of them CAT, AGT, AGTRAP, MME (reduced co-expression networks) and ATP6AP2 (enhanced co-expression networks) were also present among ACE2 seed genes. Decreased co-expression networks among top bottleneck genes were observed for EGFR, FN1, TP53, FBXO6, RNF2, ELAVL1, PCNA, and HSP90AA1. While the strongest increase in co-expression network was observed for bottleneck genes NTRK, COPS6, RAD51, PTEN, PSMA3, FRMD5, TRIM25 and APP.

#### 3.3.2. Identification of the most altered connections between ACE2-related bottleneck genes

We also observed an enhancement of the coexpression network for EWSR1. Moreover, most enhanced interactions were between HSP90AA1-MAST2 and ISYNA1(ACE2 interactor) with TRIM25. The most diminished interactions were between APP-MAST2 and EGFR-MAST2.

#### 3.3.3. Identification of the most altered connections between ACE2-related seed genes and bottleneck genes

Analysis of interactions between the seed genes and bottleneck genes showed the strongest alterations in the connection between CALM1 and RNF2 involved in cardiac development 9Figure 4 B). Also, analysis of the alterations connections between seed genes showed affected ACE2-CALM1 interaction (RankDelta=60, Rank X=155) and AGT-MME (Rank Delta=85, RankX=63). ACE2 placed in terms of Rank Delta on place 115, and Rank X 257 from 7844. In total, we identified 34 from our 69 seed genes showing changes in-expression networks. Fourteen of them were co-expressed with each other (Figure 5A)

**Figure 4.**
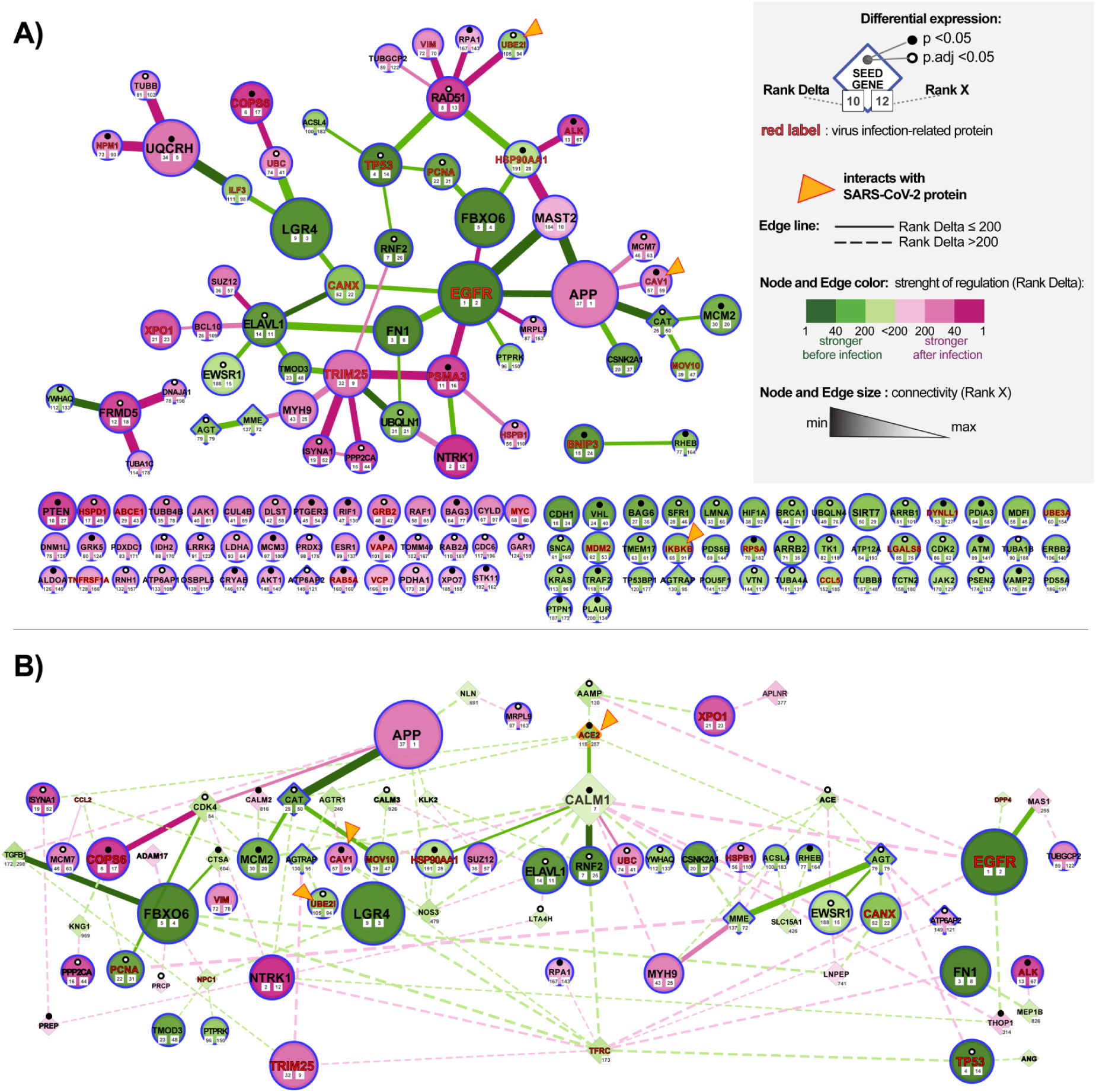
Alteration of ACE-2 networks in cardiomyocytes infected with SARS-CoV-2. (A) ACE-2 related bottleneck genes obtained by analyzing expression data of pluripotent stem cell-derived cardiomyocytes (hiPSC-CMs) after 72 hours of infection with SARS-CoV2. The network was constructed by using the NERI algorithm which integrates protein-protein interaction (PPI) Biogrid network with gene co-expression network. For clarity, we selected top genes and edges which had Rank Delta and Rank S score higher than 200; additionally, we showed nodes with the highest NERI scores which didn’t have associated edges with high scores. Genes marked with orange triangles showed direct interaction with SARS-Cov-2 [39]. Notice that EGFR and APP showed the strongest alterations in their co-expression networks. (B) PPI network between top coexpressed bottleneck genes from panel A (circular shapes) and seed genes (diamond shapes) related to the complete ACE2 network identified by us using data mining. The size of the nodes and weight of the edges is associated with Rank X score, related to biological importance, while color is associated with Rank Delta, related to the difference in co-expression network between control and disease. Notice that ACE2 shows a reduced number of connections, consistent with its initial downregulation in the early stage of infection; in later stages of infection, we can expect an inversion of observed regulation caused by the virus-relatedACE2 overexpression [40].

**Figure 5.**
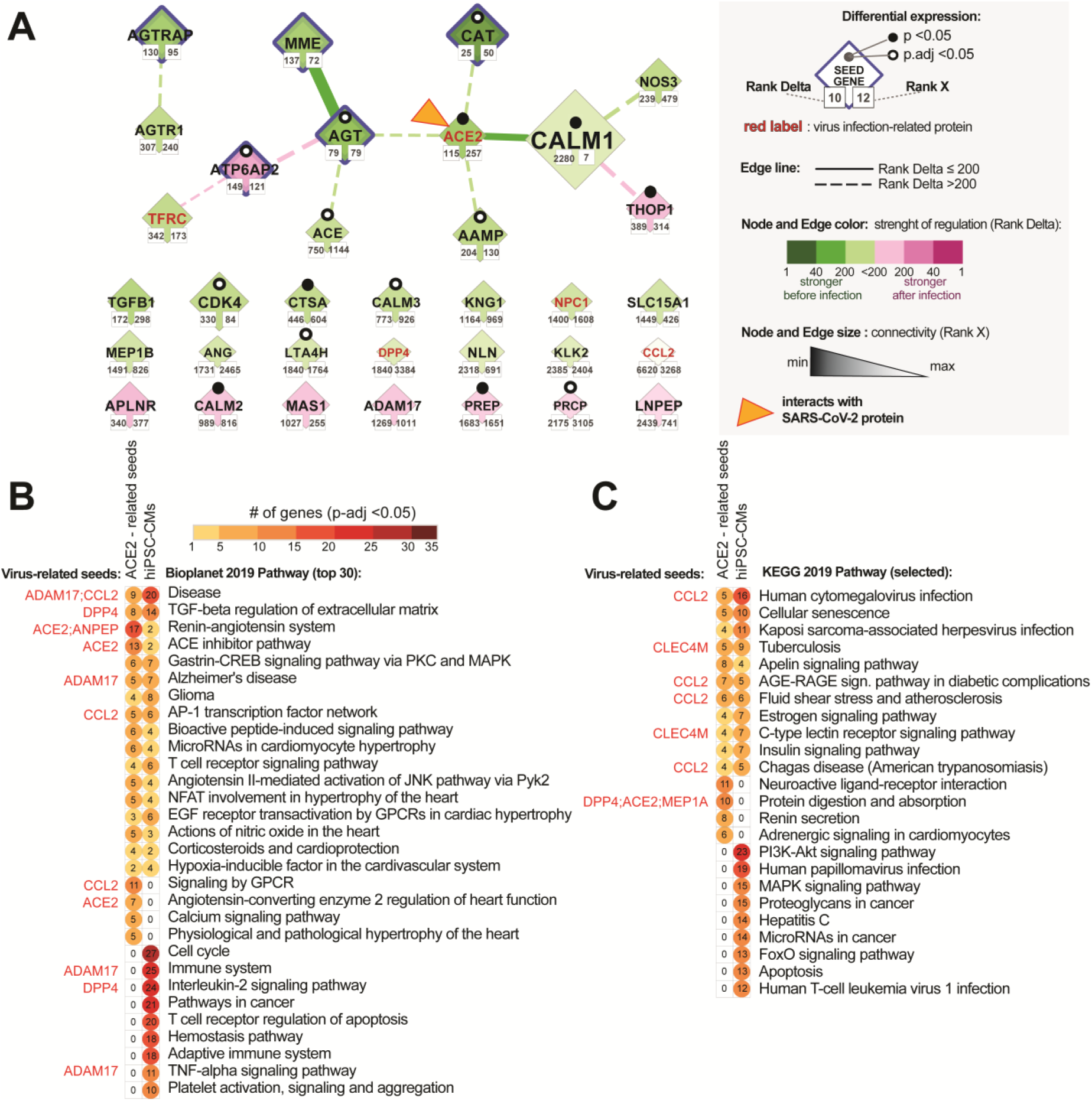
ACE2 network in infected hiPSC-CMs and top signalling pathways enriched in ACE2 interaction networks. (A) ACE2 related genes identified using data mining and co-expression network in stem cell-derived cardiomyocytes (hiPSC-CMs) 72 hours after infection. Pathway enrichment analysis of the complete ACE2 network (68 genes) and 139 top bottleneck genes identified in ACE2 related co-expression network analysis for hiPSC-CMs. Notice that the strongest altered interaction was between AGT-MME and ACE2 and CALM1, and the strongest affected nodes are AGT and CAT. (B) Bioplanet database (top 30 pathways) and (C) KEGG database (additional pathways not present in Bioplanet). Virus-infection related proteins from the complete ACE2 network are marked with red font. All circles presented on the graph are associated with significantly enriched pathways (FDR corrected p-value <0.05). In this analysis, we also included ACE2-sub-networks for heart, lungs, and nervous system, but due to high similarity with results for the complete ACE2 network, we excluded them from the figure for better clarity.

#### 3.3.4. Identification of the most altered connections between ACE2-related seed genes

The strongest interaction between ACE2 and other genes was for seed gene CALM1 (Rank Delta=2280, Rank X=155). Generally, seeds were not expected to have high-Rank X scores unless they were not a hub of interactions for other seed genes

### 3.4. Enrichment analysis of the signalling within ACE2-tissue-specific network

In order to compare our in silico predictions of the ACE2 network with its co-expression changes in infected cardiomyocytes, we performed enrichment analysis of signalling pathways using the EnrichR website (Fig 5). The aim was also to correlate those results with later disease predictions. We used 68 genes from the complete ACE-network and its subnetworks in the heart, lungs, and nervous system, as well as the top 139 bottleneck genes identified by the NERI algorithm as most affected by the SARS-Cov-2 infection. Enrichment analysis of those networks showed multiple shared pathways associated with disease-related signalling, TGF-beta regulation of extracellular matrix, renin-angiotensin pathway, Alzheimer-disease, AP-1 transcription factor network. Among top pathways shared between complete ACE2 networks, subnetworks and ACE2 network in cardiomyocytes, we observed many terms directly associated with heart functions for example microRNAs in cardiomyocyte hypertrophy, EGF receptor transactivation by GPCRs in cardiac hypertrophy, actions of nitric oxide in the heart, corticosteroids and cardioprotection. We also observed significant enrichment in all analyzed datasets of cellular senescence, apelin signalling, AGE-RAGE signalling pathway in diabetic complications, and estrogen signalling pathway identified using the KEGG database.

By comparing the complete ACE2 network to the co-expression in the hiPSC-CMs network, we observed in the former one a higher number of genes associated with renin-angiotensin related signalling, ACE inhibitor pathway, renin secretion, protein digestion, and absorption. On the other hand, in infected cardiomyocytes, there are more genes related to cell cycle signalling, interleukin-2 signalling, cancer-related pathways, T cell receptor regulation of apoptosis, and hemostasis pathway. Additionally, the platelet-degranulation pathway was enriched in infected cardiomyocytes, but also in heart and lung subnetworks, but not in the complete ACE2 network.

### 3.5. Enrichment analysis of the disease terms associated with ACE2-tissue-specific network

In order to identify the disease traits which would be helpful in precise identification of the risk groups of patients with COVID-19, we performed an enrichment analysis of the DisGenet disease and Rare Diseases AutoRIF database using the EnrichR website to evaluate phenotypes associated with ACE2 interaction in different tissues. This analysis guides the identification of phenotypes that can be triggered by ACE2-network alterations in selected tissues heart, lungs, nervous system, hiPSC-CMs as well as virus-protein-related network and complete ACE2 network. Moreover, the analysis of rare traits, usually related to single genes enabled us to precisely characterize the consequences of alterations in the specific genes from the ACE2-network.

#### 3.5.1. Analysis of the ACE2-related common diseases

The analysis of non-cancerous diseases in the DisGenet database revealed that the highest number of genes from all analyzed networks was associated with the following disease phenotypes (in the decreasing order): numerous cancerous diseases, obesity, hypertensive disease, non-insulin-dependent DM, congestive HF, coronary artery disease and atherosclerosis and were observed in all analyzed networks (Figure 6A). Enriched terms not enriched in the virus-related network, but containing virus-infection related genes were: Alzheimer’s disease, heart failure, diabetes mellitus, asthma and rheumatoid arthritis. Alzheimer’s disease and leukemia were the strongest enriched term in infected cardiomyocytes.

**Figure 6.**
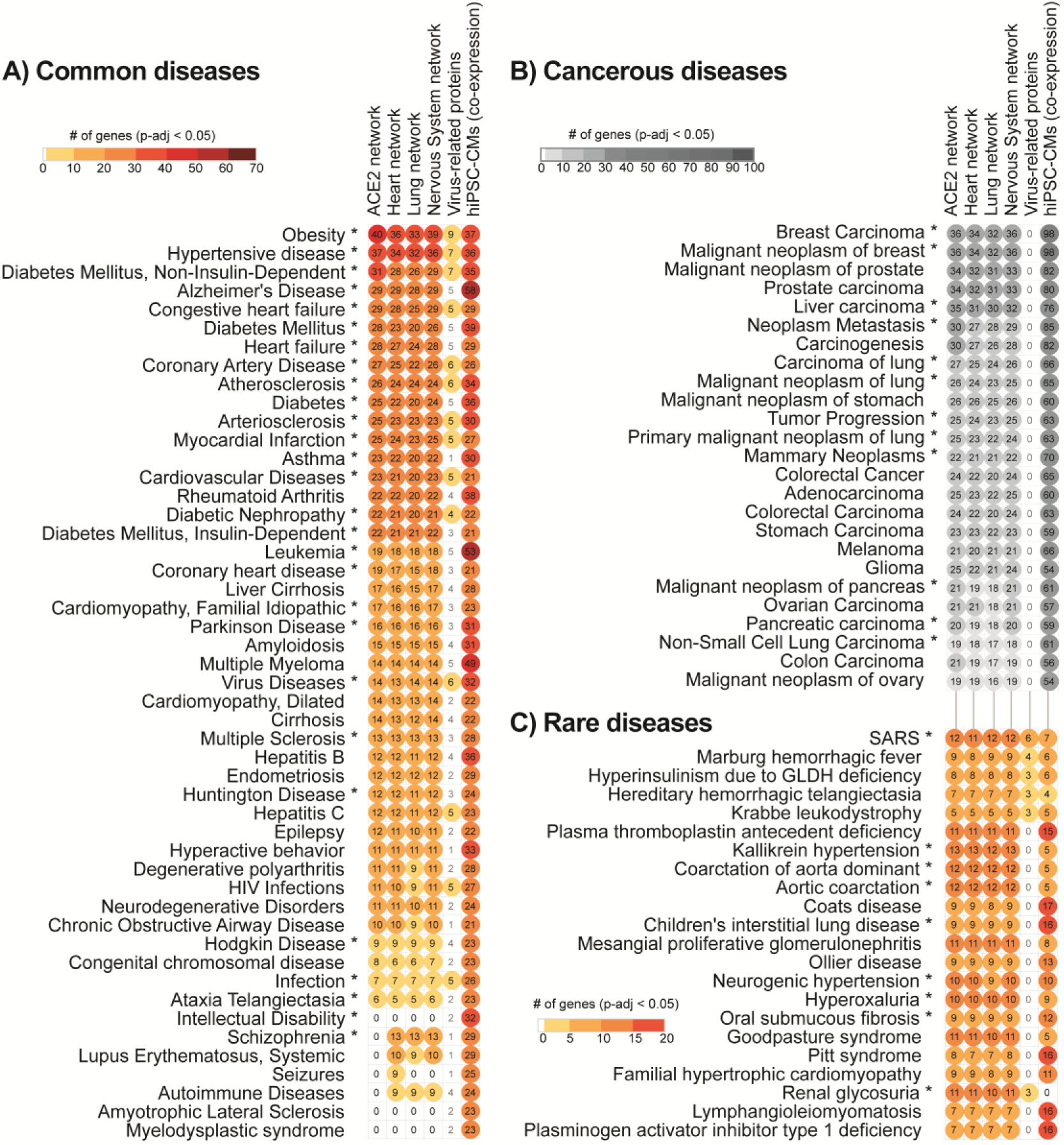
Top potential COVID-19risk groups are significantly associated with ACE2 interaction networks. Risk groups are characterized (A) common, (B) cancerous and (C) rare diseases. This list is based on enrichment analysis of DisGeNET, analyzed through the EnrichR database. We performed disease enrichment analysis in the complete ACE2 network (left-most column of symbols), subsets of this network expressed in the heart, lung, and nervous system; we performed this same analysis also for 11 virus-infection related proteins (right-most column) and ACE2-coexpression network in stem cell-derived cardiomyocytes (hiPSC-CMs) after 72 hours of infection. Diseases marked with asterisks include the ACE2 gene. For heart, lung and nervous system tissue, we used the cutoff of expression confidence >2, obtained from the Tissue2.0 database. All circles presented on the graph are associated with significantly enriched disease terms (FDR corrected p-value <0.05). Top cancer-related diseases are shown on panel B and were subset from the Common diseases by using cancer-related keywords. Notice that top diseases are already known as major risk groups in COVID-19.

#### 3.5.2. Analysis of the ACE2-related cancerous diseases

Cancerous diseases were suggested for the separate graph by using cancer-related key-words and showed the strongest enrichment of breast cancers and prostate-related cancers as well as general carcinogenesis-related processes including neoplasms metastasis (Figure 6B)

#### 3.5.2. Analysis of the ACE2-related rare diseases

The analysis focused on rare diseases revealed that the most significant ones were SARS, blood-coagulation-related diseases (Marburg hemorrhagic fever, hereditary hemorrhagic telangiectasia, plasma thromboplastin deficiency, Coats disease), and multiple diseases associated with hypertension (Kallikrein hypertension and aortic coarctation) (Figure 6C). Among rare diseases, we also observed significant enrichment of Eclampsia (average 7.1 genes from each dataset), HELLP syndrome (average 6.3 genes from each dataset) and Kawasaki disease (average 3.3 genes from each dataset) observed occasionally in COVID-19 [41].

### 3.6. Integration of ACE2 network with SARS-CoV-2/Human Interactome

To identify how the ACE2 network is connected with the SARS-CoV-2/Human Interactome and how it could affect the heart, we combined the previously published SARS-CoV-2 interactome [39] with our complete ACE2 network and top findings from the co-expression network analysis in hiPSC-CMs 72 hours after infection.

#### 3.6.1. Identification of ACE2-related genes interacting with the virus proteins

We found that three proteins from our network, ACE2, CLEC4M, and TMPRSS2, are directly interacting with virus glycoprotein S. Also top bottleneck nodes identified in NERI analysis CAV1, UBE2I, and IKBKB showed connections with virus proteins ORF3a, N and M. In total, 45 proteins from the complete ACE2 network are interacting with 38 from 94 human host proteins for SARS-CoV-2.

#### 3.6.2. Identification of ACE2-related genes interacting with the host proteins

We also found that SARS-CoV-2 interactome strongly connects with 23 bottleneck genes which were also the strongest affected in hiPSC-CMs co-expression network analysis including EGFR, APP, FN1, TP53. SARS-CoV-2 interactome strongly connects with the complete ACE2 network through INS, CDK4, CCL2, and ALB, all of them associated with atherosclerosis processes. The strongest connection between the complete ACE2 network and SARS-CoV-2/Human Interactome was with INS, which interacted with 12 host proteins from SARS-CoV-2/Human Interactome. Other top interactors were seed genes CAT, CCL2, CDK4, CALM1 connected with at least 6 host proteins. The strongest interaction for ACE2-related co-expression networks in hiPSC-CMs was between HSP90AA1 and TP53 connected with 16 host proteins from SARS interactome. Among host proteins directly interacting with virus proteins, the strongest connection with the ACE2 network occurs by ALB (30 interactors form ACE2 network) and CAV1 (13 interactors) which had affected the co-expression network after virus infection in hiPSC-CMs (Figure 7).

**Figure 7.**
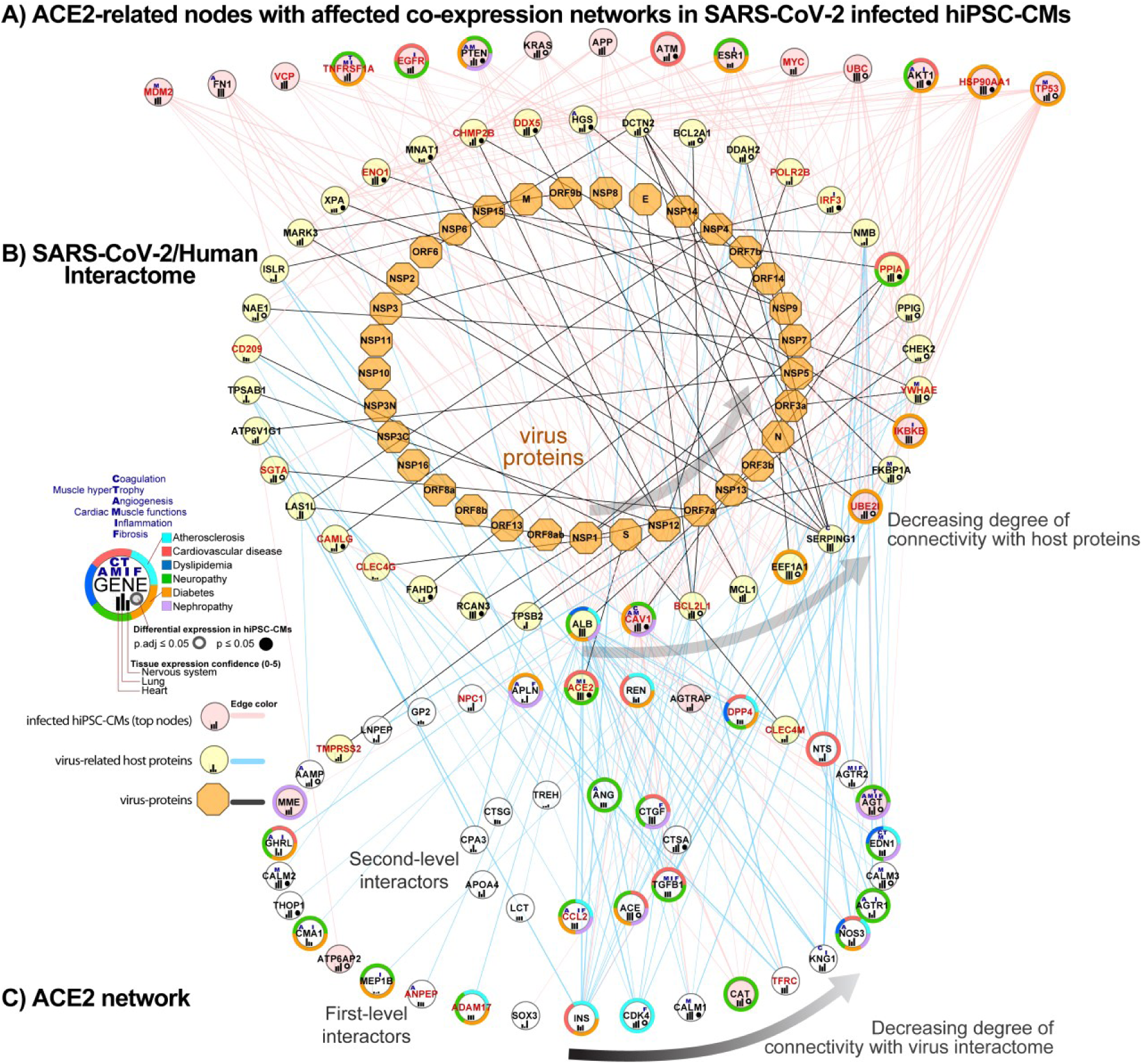
Combined ACE2 network with SARS-CoV-2/Human interactome and co-expression network in infected hiPSC-CMs. (A) Nodes identified in the co-expression network analysis of infected hiPSC-CMs using the NERI algorithm and showing the highest connectivity with SARS-CoV-2/Human interactome (top 15 genes). (B) SARS-CoV-2/Human interactome as shown in previously published work [39]. (C) ACE2 network components which interact with SARS-CoV-2/Human interactome proteins. Nodes from network A which have the highest connectivity are shown are sorted from right to left. Nodes from the networks B and C are circularly sorted by the number of connections with virus interactome. Virus proteins are shown as orange octagons, while virus-infection related human proteins have red labels.

### 3.7. ACE2 network-related miRNA predictions

In order to identify miRNAs which could play a regulatory role in COVID19, and especially its cardiovascular consequences, we performed miRNA-target predictions using the following sets of genes: 69 genes from the complete ACE2 network, gene lists from its subnetworks for heart, lungs, nervous system, virus-related proteins, as well as top bottleneck genes from the ACE2 co-expression network in hiPSC-CMs. Because among top nodes identified in the co-expression analysis of hiPSC-CMs were present only four seed genes, we decided to include for target predictions also ACE-related seed genes which showed interactions with the top bottleneck genes. For improving the precision of predictions, we analyzed miRNAs that showed expression in blood, serum, or plasma according to the Tissues2.0 database. In further analyses, we focused on miRNAs which regulated the highest number of the genes from the network as well including ACE2. Analysis of the top 10 miRNAs regulating each from those 7 networks revealed overall 10 miRNAs regulating also ACE2 gene (Figure 8a). Seven of them (hsa-miR-302c-5p, hsa-miR-27a-3p, hsa-miR-1305, hsa-miR-587, hsa-miR-26b-5p, hsa-miR-10b-5p, hsa-miR-200b-3p) were also regulating high number of virus-related proteins (Figure 8B). Among miRNAs shared across analyzed networks that were not regulating ACE2 but were present among the top 10 in all analyzed datasets, we identified *i*.*a*.hsa-miR-124-3p, hsa-miR-34a-5p, hsa-miR-548c-3p and hsa-miR-16-5p.

**Figure 8.**
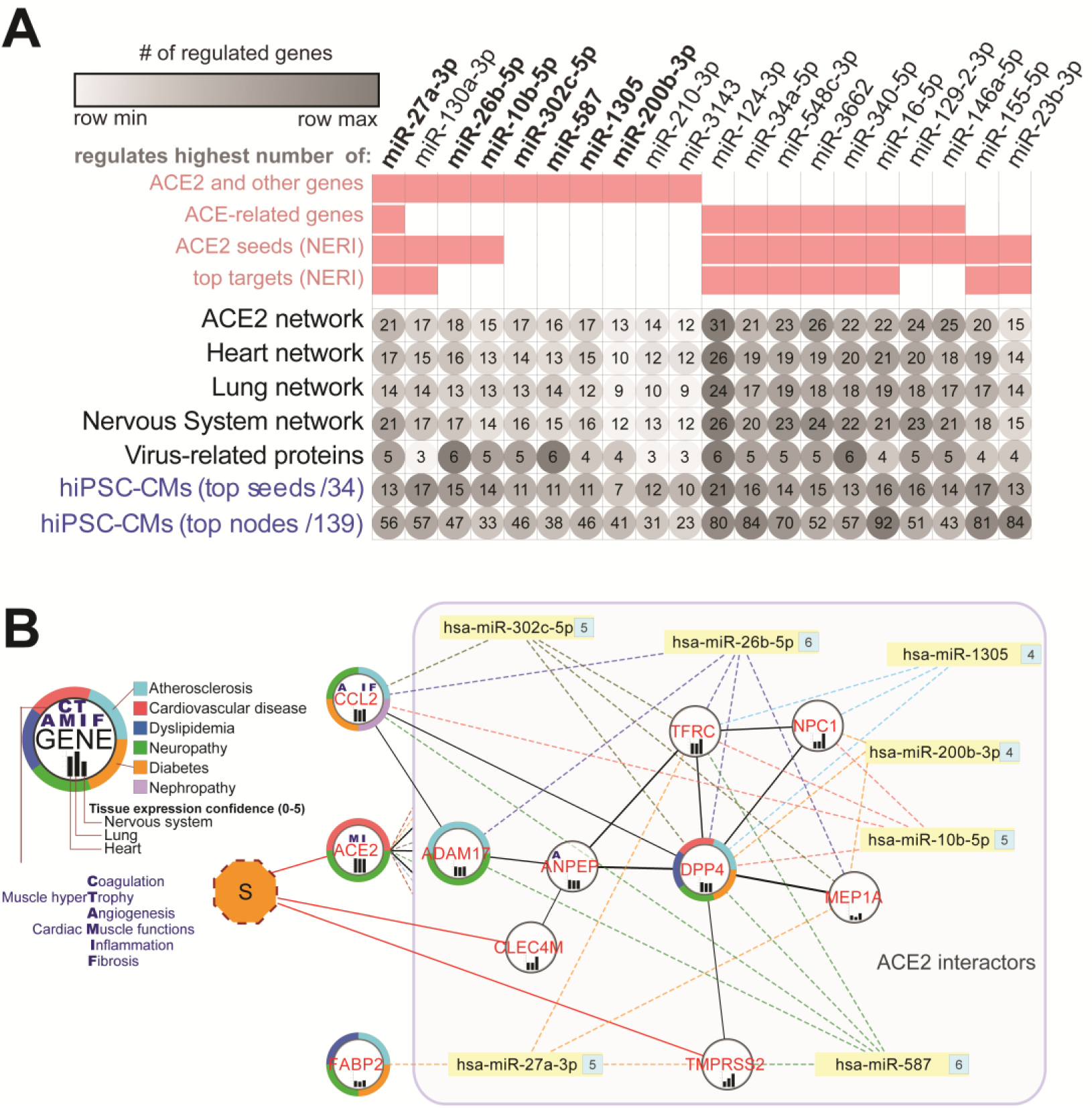
Top 20 potential miRNA modulators of the ACE2 network in COVID-19. (A) Top miRNAs regulating the highest number of genes within all ACE2 related-networks. Pink squares are showing if miRNA was present among the top 10 miRNAs in a given dataset. (B) Interaction network between virus-infection related proteins (red labels) and top miRNAs regulating ACE2 and shared between analyzed networks and regulating at least 4 virus-related proteins from the complete network. Numbers on the right side of the miRNAs depict the number of targeted genes within the network. CCL2 and FABP2 genes are not direct interactors of the ACE2, so they are presented outside of the ACE2-interactors box. “S” refers to SARS-CoV-2 S spike glycoprotein.

## 4. Discussion

In the current study, we characterized the interaction between SARS-CoV-2 infection and ACE2 functional networks with a focus on CVD. Using an integrative multi-omic approach, we described the ACE2 interaction network and evaluated its expression using the hiPSC-CMs dataset. This approach of integrating multiple sources of biological data used in our study is becoming increasingly popular [42]. There are currently numerous databases integrating, and systematizing data from different levels of biological regulation making this knowledge easily accessible. To improve the identification and prioritization of genes associated with complex diseases, some works began to integrate PPI networks with information derived from other omics data, which have contributed to a better understanding of gene functions, interactions, and pathways [25,43][44]. The integration of PPI networks and gene expression data has improved disease classification and identification of disease-specific deregulated pathways also in COVID19 [45][46][47].

The main findings of our analysis are the following: (1) Expression of ACE2 is similar in lungs and heart, which provides a rationale why the cardiovascular system is also a target of SARS-CoV-2 infection; (2) Our analysis of co-expression networks in infected hiPSC-CMs identified multiple genes and interactions associated with cardiovascular risk factors;(3) SARS-CoV-2 binding to the ACE2 receptor leads to major disturbances in signalling pathways linked to cardiac adverse outcomes in COVID19; (4) Analysis of ACE2 interaction networks revealed its association with numerous diseases including main COVID19 co-morbidities. (5) Analysis of the SARS-CoV-2 interactome revealed extensive connections with the with the top regulators of the ACE2 network; (6) We identified multiple miRNAs regulating ACE2 network (hsa-miR-302c-5p, hsa-miR-27a-3p, hsa-miR-1305, hsa-miR-587, hsa-miR-26b-5p, hsa-miR-10b-5p, hsa-miR-200b-3p; hsa-miR-124-3p and hsa-miR-16-5p).

Our hypothesis states that major implications of COVID19 cardiovascular outcomes are related to alteration in ACE2 receptor signalling associated with SARS-CoV-2 binding. Due to the difference between tissue-specific ACE2 networks, different pathological processes can be triggered in different organs. Our comparative analysis of multiple tissues showed consistency in crucial pathways including RAS signalling. This could help explain cardiac abnormalities presented in COVID-19 patients, including elevated troponin, myocarditis, arrhythmias, and sudden cardiac death [48]. Moreover, as the myocardial interaction network of ACE2 and its own expression is altered in patients with coexisting CVD, SARS-CoV-2 infection may result in greater damage to cardiomyocytes and account for greater disease acuity and poorer survival. This is consistent with recent data regarding the analysis of RNA-seq data of SARS-Cov-2 infected patients alongside controls: ACE2 network alteration seems to be the source of most recognized manifestations of COVID19 disease [40].

### 4.1. The high expression of ACE2 in the cardiovascular system explains why SARS-CoV-2 infection may target the heart

The analysis focused on the ACE2 tissue-specific expression showed that lungs and respiratory systems have similar ACE2 and TMPRSS2 expressions as in heart and cardiovascular systems and nervous systems. Recent publications regarding the impact of SARS-CoV-2 on the heart tissue showed that indeed cardiomyocytes can be infected with the virus [13][14].

### 4.2. The high expression of ACE2 in the reproductive system and endocrine glands and enrichment of cancerous diseases related to those organs suggest wider implications of SARS-CoV-2

In our analyses, we also observed strong confidence in ACE2 expression in urogenital and endocrine tissues. Moreover, we found in the genitourinary tract the highest expression of TMPRSS2 gene that is known to positively regulate viral entry into the host cell via proteolytic cleavage of ACE2. In fact, there are studies supporting the high expression of ACE2 in the urogenital tissues [49][50][51] and endocrine tissues [52], additionally, there are increasing evidence of symptoms, like male infertility [53][54], urinary tract inflammation linked to viral cystitis [55], and transmission of the virus through urine [56]. On the other hand, so far qPCR experiments to detect SARS-CoV-2 in semen and testicular biopsy didn’t find signs of virus infection [50]. We hypothesize that in this case disturbances in urogenital tissues could be associated with alternative regulation of the ACE2 network. Additionally, our analyses using the NERI algorithm showed strong alterations in cancer-related TP53, NTRK, COPS6, and RAD51 co-expression network and exceptionally high enrichment of cancerous processes associated with reproductive organs. Thus, our prediction is that additional impacts of SARS-CoV-2 infection can be observed much later than acute infection. Interestingly, TP53 is considered a master regulator of maintenance of cardiac tissue transcriptomic homeostasis [57].

### 4.2. ACE, REN, INS, KNG1, and AGT play role in cardiovascular risk factors related to SARS-CoV-2 binding to the ACE2 receptor

In our study ACE, REN, INS, KNG1, and AGT showed the highest connectivity within the complete ACE2 network and association with enriched diseases that are known to be a risk factor for a severe course of COVID-19. Our analysis of the data related to infected hiPSC-CMs confirmed changes in co-expression networks of ACE, AGT (also present among bottleneck genes), and KNG1. Both ACE and AGT also showed differential expressions in hiPSC-CMs. On the other hand, expression levels of INS, and REN were very low thus suggesting rather their role in other than cardiac tissues. ACE and ACE2interplay with the RAS pathway and plasma kallikrein-kinin system (KKS) which component is KNG1, a hormonal pathway that modulates the intrinsic blood coagulation system, angiogenesis, the complement pathway and bradykinin-related inflammation. Deregulation of KKS may result in thromboembolic complications and lead to sepsis exacerbation in infections [58–60]. Deregulation of AGT which is a crucial component of the RAS pathway is associated with the pathogenesis of essential hypertension and atrial fibrillation.

### 4.3. Analysis of infected hiPSC-CMs using the NERI algorithm revealed affected co-expression networks ofEGFR, APP, and CALM1 implicating their role in cardiac and thromboembolic complications

EGFR loss in vascular smooth muscle cells and cardiomyocytes leads to arterial hypotension and cardiac hypertrophy [61]. EGFR was found as a pivotal regulator of thrombin-mediated inflammation. It also plays a role in changes from lethal versus non-lethal influenza infections [56,62]. EGFR showed decreased signalling in infected hiPSC-CMs.

APP is a precursor protein for Amyloid-beta (Aβ) peptide which is massively released from the blood to nearby tissue upon the activation of platelets and has strong antibiotic activity against viruses, bacteria, and fungi [63]. Accumulation of Aβ in tissues is observed in Alzheimer’s disease, glaucoma, cancerous disease, myocardium with diastolic dysfunction, and the placenta during preeclampsia [64,65]. All those diseases were enriched in our study. APP showed an increased co-expression network in infected hiPSC-CMs. Identification of this interplay between EGFR and APP can play an important role in intervention targets in COVID19 treatment.

The analysis focused on close ACE2 interactors revealed its strongly affected connection with CALM1 in infected hiPSC-CMs. CALM1 regulates the function of ion channels playing a role in platelet aggregation and cardiomyocytes activity. It is also an important ACE2 interactor playing a role in viral pathogenesis [66,67]. In our analysis, CALM1 showed interactions also with bottleneck node PTEN affecting cardiomyocytes contractions and growth.

### 4.4. Alterations in the ACE2 interaction network can aggravate major comorbidities in COVID19 through related signalling pathways

Our analysis of ACE2 interaction networks including co-expression networks in infected cardiomyocytes showed that change in ACE2 receptor activity can lead to significant disturbances in signalling pathways linked to well-known complications in COVID-19. Those pathways included TGF-beta regulation of extracellular matrix, renin-angiotensin system AP-1 transcription factor network, apelin signalling, AGE-RAGE signalling pathway in diabetic complications, and estrogen signalling Signalling pathways the most affected in hiPSC-CMs were related to cell cycle (i.a. Pathways in cancer, PI3K-Akt signalling pathway), immune system (i.a. interleukin-2 signalling pathway, T-cell receptor regulation of apoptosis), hemostasis and platelet activation. Those results were supported by our subsequent analysis of disease-related phenotypes which are the major comorbidities in COVID19 cancerous diseases, obesity, hypertensive disease, diabetes, and Alzheimer’s disease.

### 4.5. Renin-angiotensin pathway, AGE-RAGE, and Apelin signalling as fundamental mediators of the blood pressure dysregulation mediated through ACE2 in COVID-19

In our *in silico* analysis using ACE2 functional networks, we found that RAS/ACE2, AGE-RAGE, and Apelin signalling pathways play an important role in SARS-CoV-2 infection. These pathways have a crucial role in the pathogenesis of DM, CVD, and blood pressure regulations. Abnormalities of ACE2/RAS pathway signalling and deregulation of angiotensin II as a fundamental mediator of this axis are closely related to the pathophysiology of hypertension and progression of cardiovascular remodeling [68] [69]. Therefore, binding of the SARS-CoV-2 to the ACE2 receptor leading to disturbances in the pathways of these key regulators might explain the adverse outcome in COVID-19 patients with the coexistence of the above mentioned clinical conditions. Also, apelin signalling is involved in many physiological processes such as energy metabolism, blood pressure regulation, and cardiac contractility and plays an important role in organ and tissue pathologies including, DM, obesity, HF as well as HIV-1 infection [70]. After the virus enters the cells, ACE2 is likely to decrease its activity, thus favoring an increase of the ACE/ACE2 balance toward the prevalence of ACE arm in the RAS which causes an increase of ROS production, vasoconstriction, and inflammation [71]. On the other hand, a recent COVID-19 related study showed decreased expression of ACE in combination with increases in ACE2 likely causing bradykinin-related increases in vascular dilation, vascular permeability and hypotension explaining many of the symptoms being observed in COVID-19 [40].

### 4.6. The role of virus-infection related proteins from ACE2 network in COVID-19 adverse outcomes

Analysis of the connection between SARS-CoV-2 interactome revealed direct interaction of the virus glycoprotein S with three of 11 virus-infection related proteins identified incomplete ACE2 network (ACE2, CLEC4M, and TMPRSS2). We also identified DE bottleneck nodes (CAV1, UBE2I) interacting with other virus proteins. Interestingly, CAV1 was identified as a possible alternative receptor for SARS-CoV and Canine respiratory coronavirus (CRCoV), which may be associated with the virus infection, replication, assembly, and budding [72][72,73]. CAV1 showed an enhanced co-expression network. Additionally, we observed multiple connections between host genes from virus interactome and most affected genes in hiPSC-CMs. Those genes included TP53, HSP90AA1 but also ESR1, FN1, APP and EGFR and seed genes CAT, AGT, AGTRAP, DPP, CCL2 and MME. The especially interesting genes are discussed below.

HSP90AA1 was recently shown in pre-print as reducing SARS-CoV-2 viral replication, and TNF and IL1B mRNA levels [74]. Additionally, in our analysis, the strongest affected interaction in cardiomyocytes was observed between HSP90AA1 and MAST2. MAST2 regulates IL12 production in macrophages and shows association with red blood cell distribution width which was identified recently as a biomarker of COVID19 mortality [75]. Interestingly, MAST2 also showed an affected connection with the top bottleneck gene APP. HSP90AA1 and seed gene CAT showed significant differential expression and a strong reduction in co-expression networks in infected hiPSC-CMs in our study. CAT is considered the most effective catalyst for the decomposition of H2O2, regulating the production of cytokines, protecting from oxidative injury, and repressing replication of SARS-CoV-2 [76].

FN1 in our study showed decreased signalling in hiPSC-CMs. FN1 inhibition attenuates fibrosis and improves cardiac function in a model of heart failure [77]. Interaction of human plasma fibronectin with viral proteins of HIV suggests that it’s binding to virus particles may reduce viremia and thus may be involved in the clearance of viral proteins from the cells [78].

Higher plasma DPP4 can be found among patients with obesity, metabolic syndrome, and DM, who are at risk of a severe course of COVID-19 [79]. DPP4 knock-in mice were found more susceptible to MERS-CoV infections which resulted in the severe inflammatory response and lethal lung disease.[80,81] Therefore, it should be further investigated whether DPP4 inhibitors, widely used for the treatment of DM, may act as therapeutic drugs for ARDS caused by SARS-CoV-2 infection.[82] Our study revealed the possible interaction between DPP and EGFR which was identified as the most affected gene in co-expression network analysis.

Among affected virus-related-infection proteins, we also identified CCL2. CCL2 protein has been implicated in lung inflammatory disorders and contributes to the development of pulmonary fibrosis [23,83]. It is worth mentioning that among SARS-CoV-infected patients the level of pro-inflammatory cytokines, especially CCL2 and TGF-β1 (both affected in infected hiPSC-CMs) were increased in cells expressing ACE2, while this could not be seen in tissue with undetectable ACE2 expression [84]. A comparable pattern of inflammatory cytokines was found in SARS-CoV-2 infection as well [85]. This makes CCL2 a promising link between ACE2 and cytokine storm associated with severe COVID-19 disease.

Altogether, our results suggest that not only ACE2 is affected by the entrance of the virus to the cardiomyocytes, but that this virus also affects multiple of ACE-2 interactors and that the ACE-2 network can be as well a part of virus propagation machinery.

### 4.7 miRNAs as promising antiviral modulators of the ACE2 network and a potential biomarker of HF associated with COVID-19

In the initial step of our study we identified miRNAs that may regulate the expression of ACE2 networks and related processes. To our best knowledge, we present here a novel results on a potential role of miRNAs as a diagnostic and prognostic tool in heart muscle injury in the course of SARS-CoV-2 infection.

#### miR-1305 and miR-587: TGF-β signalling pathway regulators in HF progression

MiR-1305 and miR-587 were found to regulate the expression of TGF-β pathway members related to virus infections and lymphocytes T activation, SMAD3, and SMAD4 [86,87][88][89], ventricular remodeling, myocardial fibrosis and hypertrophy and, as a result, HF progression [90]. The highest expression of miR-587 was found in platelets of patients with acute coronary syndrome and was closely related to the severity of coronary artery stenosis [91].

#### miR-26b-5p: anti-fibrotic agent and AGTR1-dependent hypertension modulator

In our study, we found that miR-26b-5p may play an important role in the pathogenesis of HF in COVID-19 patients. Noteworthy, literature data suggest that AGTR1 can modulate hypertension, via the regulation of miR-26b-5p in arachidonic acid metabolism. Additionally, miR-26b-5p has an anti-fibrotic effect in the liver, in the diabetic mouse myocardium, and in Ang-II-induced mouse cardiac fibroblasts [92].

#### miR-302c-5p: potential antiviral therapeutic and biomarker of HF

Another miRNA from our network affecting ACE2 was miR-302c-5p playing an important role in many viral infections [93–95]. A study reported an association between the miR-302, cytokine storm and showed the potential of miR-302 as an antiviral therapeutic [95]. Our bioinformatic analysis for the first time showed the importance of miR-302c-5p in SARS-CoV-2 infection. Apart from the crucial function of miR-302 in viral infections, it may be also associated with CVD [96,97], as circulating miR-302 was positively correlated with NT-proBNP levels in acute HF patients and showed strong potential as a novel biomarker for the diagnosis and the differentiation of disease severity of acute HF [96].

#### miR-27a-3p: a potential biomarker of acute HF and NF-κB signalling regulator

Also, miR-27a-3p targeting ACE2 and other genes from its network in our study were found to be involved in the inflammatory response and oxidative stress through several pathways including PPAR-γ, NF-κB, and PI3K/AKT/Nrf2 signalling [98–100]. In the animal model of acute lung injury, expression of miR-27a-3p in alveolar macrophages was significantly decreased, while overexpression of miR-27a-3p suppressed NF-κB activation and alleviated acute lung injury by binding to its target NFKB1 [98]. Moreover, it was also found that miR-27a-3p may target pathways related to atherosclerosis, and may act as a potential biomarker of acute HF [101,102].

#### hsa-miR-16-5p: modulates inflammatory signalling and cytokines including IL-1β, IL-6, and TNF-α, NF-κB mTOR-related pathways

*hsa-miR-16-5p* was found to affect a phenotypic change of T cells, modulate inflammatory signalling and cytokines including IL-1β, IL-6 and TNF-α, NF-κB mTOR-related pathways and genes [103,104]. Additionally, MiR-16-5p has been linked with the pathogenesis of several infectious diseases such as HIV-1 infection and malaria [105–107]. It is worth mentioning that miR-16-5p as a plasma diagnostic biomarker is able to distinguish severe and mild viral infections [108] and early HIV-1 infection from healthy individuals [106,109].

#### hsa-miR-124-3p: has potentially an aggravating role in cardiovascular consequences of COVID19

MiR-124-3p was identified by us as regulating the highest number of genes in hiPSC-CMs. Literature data show its aggravating role in failing hearts by suppressing CD151-facilitated angiogenesis in the heart [110]. miR-124-3p dysregulates NSC maintenance through repression of the transferrin receptor (TFRC) in Zika virus infection [111]. TFRC is linked with cardiomyopathy and was identified in our study as closely interacting with ACE2 in infected cardiomyocytes and strongly connected with SARS interactome.

## 5. Conclusions

This comprehensive analysis provides novel information regarding the complexity of signalling pathways of SARS-CoV-2 infection affecting the cardiovascular system with a focus on cardiomyocytes and forms a basis for the creation of predictive tools and introduction of therapy to improve outcome in COVID-19, and therefore has a potential to reduce economic consequences of the global pandemic. We believe that the results of our analysis could be further validated in laboratory and clinical settings and help to create a paradigm for future studies in this field. MiRNAs identified for the first time in this study can serve as potential biomarkers helping with the identification of the pathological changes in COVID-19 or serve as therapeutic targets due to their stability in the serum, forming a basis for personalized therapy.

## Supplementary Materials

Supp Table 1. List of GO terms used for annotation of the genes to specific processes and functions. Supp Table 2. Description of genes associated with complete ACE2 interaction network

## Author Contributions

Bioinformatic analysis: Z.W., S.N.S, D.C.M, Writing—original draft preparation, Z.W., C.E., D.J., J.M.S.-M., M.P.; writing-review and editing, Z.W., C.E., D.J., R.P., J.M.S.-M., M.P.; visualization, Z.W., C.E., R.P.; supervision, Z.W., J.M.S.-M., and M.P. The article was published as a result of the collaboration within the I-COMET team.

## Funding

This work was implemented with CEPT infrastructure financed by the European Union-the European Regional Development Fund within the Operational Program “Innovative economy” for 2007–2013.

## Acknowledgments

In this section you can acknowledge any support given which is not covered by the author contribution or funding sections. This may include administrative and technical support, or donations in kind (e.g., materials used for experiments).

## Conflicts of Interest

Authors declare no COI related to this work.

## Abbreviations

ACE2: Angiotensin-Converting Enzyme 2
ACE: Angiotensin-Converting Enzyme 1
ARDS: Acute respiratory distress syndrome
COVID-19: Coronavirus disease 2019
CVD: Cardiovascular disease
DEG: Differentially expressed genes
DM: Diabetes mellitus
eNOS: Endothelial nitric oxide synthase
GO: Gene Ontology
HF: Heart failure
HK: High molecular weight kininogen
INS: Insulin
KKS: Kallikrein-Kinin system
KNG1: Kininogen 1
MERS: Middle-East respiratory syndrome coronavirus
MI: Myocardial infarction
miRNAs, miR: MicroRNAs
NT-proBNP: N-terminal pro-B-type natriuretic peptide
PPI: Protein-protein interaction
RAS: Renin-angiotensin system
REN: Renin
ROS: Reactive oxygen species
SARS-CoV-2: Severe acute respiratory syndrome coronavirus 2
TMPRSS2: Transmembrane protease serine 2

## Gene/protein

ACE2: Angiotensin-Converting Enzyme 2
ADAM17: ADAM metallopeptidase domain 17
AGT: Angiotensinogen
AGTR1: Angiotensin II Receptor Type 1
ALB: Albumin
APP: Amyloid Beta Precursor Protein
ANPEP: Alanyl Aminopeptidase
APN/CD13: Aminopeptidase N/CD13
ATP6AP2: ATPase H+ Transporting Accessory Protein 2
CALM1: Calmodulin 1
CAT: Catalase
CAV1: Caveolin-1
CCL2: C-C Motif Chemokine Ligand 2
CCN2: Cellular Communication Network Factor 2
CDHR2: Cadherin Related Family Member 2
CDK4: Cyclin-Dependent Kinase 4
CLEC4M: C-Type Lectin Domain Family 4 Member M
CTSA: Cathepsin A
CTSG: Cathepsin G
DPP4: Dipeptidyl Peptidase 4
EGFR: Epidermal Growth Factor Receptor
ENPEP: Glutamyl Aminopeptidase
EV71: Enterovirus 71
FABP2: Fatty acid-binding protein 2
FoxO2: Forkhead box O-2
INS: Insulin
KNG1: Kininogen 1
KPNA2: Karyopherin Subunit Alpha 2
LNPEP: Leucyl And Cystinyl Aminopeptidase
LTA4H: Leukotriene A4 hydrolase
MAPK: Mitogen-activated protein kinase
MCP-1: Monocyte chemoattractant protein-1
MEP1A: Meprin A subunit alpha
MME: Membrane metalloendopeptidase
MS4A10: Membrane Spanning 4-Domains A10
NFKB1: Nuclear Factor Kappa B Subunit 1
NF-κB: Nuclear factor kappa-light-chain-enhancer of activated B cells
NPC1: Niemann-Pick disease, type C1
PAI-1: Plasminogen activator inhibitor-1
PDGFR-b: Platelet-derived growth factor receptor-beta
PPARγ: Peroxisome proliferator-activated receptor-gamma
PRCP: Prolylcarboxypeptidase
TFRC: Transferrin Receptor
TGFB1: Transforming Growth Factor Beta 1
THOP1: Thimet Oligopeptidase 1
TMPRSS2: Transmembrane protease, serine 2
VEGFR2: Vascular endothelial growth factor receptor 2

## Signalling pathways

AGE-RAGE: Advanced glycation endproducts -Receptor for Advanced glycation end products
ERK1/2/AP-1: Extracellular signal-regulated kinases 1/2/AP-1
PI3K/AKT/Nrf2: Phosphatidylinositol 3’-kinase/AKT/NF-E2-related factor 2
ERK1/2/AP-1: Extracellular signal-regulated kinases 1/2/AP-1
PI3K/AKT/Nrf2: Phosphatidylinositol 3’-kinase/AKT/NF-E2-related factor 2

## Notes

### Competing Interest Statement

The authors have declared no competing interest.

### Summary of Updates

We made substantial changes in the design of the study by including analysis of the ACE2 co-expression networks of publicly available expression data regarding infected stem cell-derived cardiomyocytes (hiPSC-CMs). This enabled us to more precisely point out the consequences of alteration of the ACE2 network in the cardiovascular system.

## References

1. Zhu, N.; Zhang, D.; Wang, W.; Li, X.; Yang, B.; Song, J.; Zhao, X.; Huang, B.; Shi, W.; Lu, R.; et al. A Novel Coronavirus from Patients with Pneumonia in China, 2019. New England Journal of Medicine 2020, 382, 727–733.

2. Wu, C.; Chen, X.; Cai, Y.; Xia, J. ‘an; Zhou, X., Xu, S.; Huang, H.; Zhang, L.; Zhou, X.; Du, C.; et al. Risk Factors Associated With Acute Respiratory Distress Syndrome and Death in Patients With Coronavirus Disease 2019 Pneumonia in Wuhan, China. JAMA Intern. Med. 2020, doi: 10.1001/jamainternmed.2020.0994.

3. Yang, J.; Zheng, Y.; Gou, X.; Pu, K.; Chen, Z.; Guo, Q.; Ji, R.; Wang, H.; Wang, Y.; Zhou, Y. Prevalence of comorbidities and its effects in coronavirus disease 2019 patients: A systematic review and meta-analysis. International Journal of Infectious Diseases 2020, 94, 91–95.

4. Li, B.; Yang, J.; Zhao, F.; Zhi, L.; Wang, X.; Liu, L.; Bi, Z.; Zhao, Y. Prevalence and impact of cardiovascular metabolic diseases on COVID-19 in China. Clinical Research in Cardiology 2020, 109, 531–538.

5. Leung, G.M.; Hedley, A.J.; Ho, L.-M.; Chau, P.; Wong, I.O.L.; Thach, T.Q.; Ghani, A.C.; Donnelly, C.A.; Fraser, C.; Riley, S.; et al. The Epidemiology of Severe Acute Respiratory Syndrome in the 2003 Hong Kong Epidemic: An Analysis of All 1755 Patients. Annals of Internal Medicine 2004, 141, 662.

6. Assiri, A.; Al-Tawfiq, J.A.; Al-Rabeeah, A.A.; Al-Rabiah, F.A.; Al-Hajjar, S.; Al-Barrak, A.; Flemban, H.; Al-Nassir, W.N.; Balkhy, H.H.; Al-Hakeem, R.F.; et al. Epidemiological, demographic, and clinical characteristics of 47 cases of Middle East respiratory syndrome coronavirus disease from Saudi Arabia: a descriptive study. Lancet Infect. Dis. 2013, 13, 752–761, doi:10.1016/S1473-3099(13)70204-4.

7. Madjid, M.; Safavi-Naeini, P.; Solomon, S.D.; Vardeny, O. Potential Effects of Coronaviruses on the Cardiovascular System. JAMA Cardiology 2020, 5, 831.

8. Wang, K.; Gheblawi, M.; Oudit, G.Y. Angiotensin Converting Enzyme 2: A Double-Edged Sword. Circulation 2020.

9. Gheblawi, M.; Wang, K.; Viveiros, A.; Nguyen, Q.; Zhong, J.-C.; Turner, A.J.; Raizada, M.K.; Grant, M.B.; Oudit, G.Y. Angiotensin-Converting Enzyme 2: SARS-CoV-2 Receptor and Regulator of the Renin-Angiotensin System. Circulation Research 2020, 126, 1456–1474.

10. Gheblawi, M.; Wang, K.; Viveiros, A.; Nguyen, Q.; Zhong, J.-C.; Turner, A.J.; Raizada, M.K.; Grant, M.B.; Oudit, G.Y. Angiotensin-Converting Enzyme 2: SARS-CoV-2 Receptor and Regulator of the Renin-Angiotensin System. Circulation Research 2020, 126, 1456–1474.

11. Patel, V.B.; Zhong, J.-C.; Grant, M.B.; Oudit, G.Y. Role of the ACE2/Angiotensin 1–7 Axis of the Renin–Angiotensin System in Heart Failure. Circulation Research 2016, 118, 1313–1326.

12. Hoffmann, M.; Kleine-Weber, H.; Krüger, N.; Müller, M.; Drosten, C.; Pöhlmann, S. The novel coronavirus 2019 (2019-nCoV) uses the SARS-coronavirus receptor ACE2 and the cellular protease TMPRSS2 for entry into target cells.

13. Sharma, A.; Garcia, G. Jr; Wang, Y., Plummer, J.T.; Morizono, K.; Arumugaswami, V.; Svendsen, C.N. Human iPSC-Derived Cardiomyocytes Are Susceptible to SARS-CoV-2 Infection. Cell Reports. Medicine 2020, 1, 100052, doi:10.1016/j.xcrm.2020.100052.

14. Bojkova, D.; Wagner, J.U.G.; Shumliakivska, M.; Aslan, G.S.; Saleem, U.; Hansen, A.; Luxán, G.; Günther, S.; Pham, M.D.; Krishnan, J.; et al. SARS-CoV-2 infects and induces cytotoxic effects in human cardiomyocytes 2020, 2020.06.01.127605.

15. Pietro Bulfamante, G.; Perrucci, G.L.; Falleni, M.; Sommariva, E.; Tosi, D.; Martinelli, C.; Songia, P.; Poggio, P.; Carugo, S.; Pompilio, G. Evidence of SARS-CoV-2 transcriptional activity in cardiomyocytes of COVID-19 patients without clinical signs of cardiac involvement. medRxiv 2020, 2020.08.24.20170175, doi: 10.1101/2020.08.24.20170175.

16. Oudit, G.Y.; Kassiri, Z.; Jiang, C.; Liu, P.P.; Poutanen, S.M.; Penninger, J.M.; Butany, J. SARS-coronavirus modulation of myocardial ACE2 expression and inflammation in patients with SARS. Eur. J. Clin. Invest. 2009, 39, 618–625, doi: 10.1111/j.1365-2362.2009.02153.x.

17. Shi, S.; Qin, M.; Shen, B.; Cai, Y.; Liu, T.; Yang, F.; Gong, W.; Liu, X.; Liang, J.; Zhao, Q.; et al. Association of Cardiac Injury With Mortality in Hospitalized Patients With COVID-19 in Wuhan, China. JAMA Cardiol 2020, doi: 10.1001/jamacardio.2020.0950.

18. Clerkin, K.J.; Fried, J.A.; Raikhelkar, J.; Sayer, G.; Griffin, J.M.; Masoumi, A.; Jain, S.S.; Burkhoff, D.; Kumaraiah, D.; Rabbani, L.; et al. Coronavirus Disease 2019 (COVID-19) and Cardiovascular Disease. Circulation 2020, doi: 10.1161/CIRCULATIONAHA.120.046941.

19. Guo, T.; Fan, Y.; Chen, M.; Wu, X.; Zhang, L.; He, T.; Wang, H.; Wan, J.; Wang, X.; Lu, Z. Cardiovascular Implications of Fatal Outcomes of Patients With Coronavirus Disease 2019 (COVID-19). JAMA Cardiol 2020, doi: 10.1001/jamacardio.2020.1017.

20. Kanehisa, M. The KEGG Database. “In Silico” Simulation of Biological Processes 91–103.

21. Doncheva, N.T.; Morris, J.H.; Gorodkin, J.; Jensen, L.J. Cytoscape StringApp: Network Analysis and Visualization of Proteomics Data. J. Proteome Res. 2019, 18, 623–632, doi: 10.1021/acs.jproteome.8b00702.

22. Zhang, H.; Penninger, J.M.; Li, Y.; Zhong, N.; Slutsky, A.S. Angiotensin-converting enzyme 2 (ACE2) as a SARS-CoV-2 receptor: molecular mechanisms and potential therapeutic target. Intensive Care Medicine 2020, 46, 586–590.

23. Chen, I.-Y.; Chang, S.C.; Wu, H.-Y.; Yu, T.-C.; Wei, W.-C.; Lin, S.; Chien, C.-L.; Chang, M.-F. Upregulation of the Chemokine (C-C Motif) Ligand 2 via a Severe Acute Respiratory Syndrome Coronavirus Spike-ACE2 Signalling Pathway. J. Virol. 2010, 84, 7703–7712, doi: 10.1128/JVI.02560-09.

24. pubmeddev; Kuan TC, E. al Identifying the regulatory element for human angiotensin-converting enzyme 2 (ACE2) expression in human cardiofibroblasts. - PubMed - NCBI Available online: https://www.ncbi.nlm.nih.gov/pubmed/21864606 (accessed on Apr 22, 2020).

25. Wicik, Z.; Jales Neto, L.H., Guzman, L.E.; Pavão, R.; Takayama, L.; Caparbo, V.; Lopes, N.; Pereira, A.C.; Pereira, R.M.R. The crosstalk between bone metabolism, lncRNAs, microRNAs and mRNAs in coronary artery calcification. Genomics 2020, doi: 10.1016/j.ygeno.2020.09.041.

26. Palasca, O.; Santos, A.; Stolte, C.; Gorodkin, J.; Jensen, L.J. TISSUES 2.0: an integrative web resource on mammalian tissue expression. Database 2018, 2018, doi: 10.1093/database/bay003.

27. Shannon, P.; Markiel, A.; Ozier, O.; Baliga, N.S.; Wang, J.T.; Ramage, D.; Amin, N.; Schwikowski, B.; Ideker, T. Cytoscape: a software environment for integrated models of biomolecular interaction networks. Genome Res. 2003, 13, 2498–2504, doi: 10.1101/gr.1239303.

28. Simões, S.N.; Martins, D.C.; Pereira, C.A.B.; Hashimoto, R.F.; Brentani, H. NERI: network-medicine based integrative approach for disease gene prioritization by relative importance. BMC Bioinformatics 2015, 16, 1–14, doi: 10.1186/1471-2105-16-S19-S9.

29. Eyileten, C.; Wicik, Z.; De Rosa, S.; Mirowska-Guzel, D.; Soplinska, A.; Indolfi, C.; Jastrzebska-Kurkowska, I.; Czlonkowska, A.; Postula, M. MicroRNAs as Diagnostic and Prognostic Biomarkers in Ischemic Stroke—A Comprehensive Review and Bioinformatic Analysis. Cells 2018, 7, 249.

30. Rani, J.; Mittal, I.; Pramanik, A.; Singh, N.; Dube, N.; Sharma, S.; Puniya, B.L.; Raghunandanan, M.V.; Mobeen, A.; Ramachandran, S. T2DiACoD: A Gene Atlas of Type 2 Diabetes Mellitus Associated Complex Disorders. Sci. Rep. 2017, 7, doi: 10.1038/s41598-017-07238-0.

31. Huang, D.W.; Sherman, B.T.; Lempicki, R.A. Bioinformatics enrichment tools: paths toward the comprehensive functional analysis of large gene lists. Nucleic Acids Res. 2009, 37, 1–13, doi: 10.1093/nar/gkn923.

32. Chen, E.Y.; Tan, C.M.; Kou, Y.; Duan, Q.; Wang, Z.; Meirelles, G.; Clark, N.R.; Ma’ayan, A. Enrichr: interactive and collaborative HTML5 gene list enrichment analysis tool. BMC Bioinformatics 2013, 14, 128.

33. Ru, Y.; Kechris, K.J.; Tabakoff, B.; Hoffman, P.; Radcliffe, R.A.; Bowler, R.; Mahaffey, S.; Rossi, S.; Calin, G.A.; Bemis, L.; et al. The multiMiR R package and database: integration of microRNA–target interactions along with their disease and drug associations. Nucleic Acids Research 2014, 42, e133–e133.

34. Pordzik, J.; Jakubik, D.; Jarosz-Popek, J.; Wicik, Z.; Eyileten, C.; De Rosa, S.; Indolfi, C.; Siller-Matula, J.M.; Czajka, P.; Postula, M. Significance of circulating microRNAs in diabetes mellitus type 2 and platelet reactivity: bioinformatic analysis and review. Cardiovascular Diabetology 2019, 18.

35. Sabatino, J.; Wicik, Z.; De Rosa, S.; Eyileten, C.; Jakubik, D.; Spaccarotella, C.; Mongiardo, A.; Postula, M.; Indolfi, C. MicroRNAs fingerprint of bicuspid aortic valve. J. Mol. Cell. Cardiol. 2019, 134, 98–106, doi: 10.1016/j.yjmcc.2019.07.001.

36. Barabási, A.-L.; Gulbahce, N.; Loscalzo, J. Network medicine: a network-based approach to human disease. Nature Reviews Genetics 2011, 12, 56–68.

37. Eley, B. Faculty Opinions recommendation of Neurological associations of COVID-19. Faculty Opinions – Post-Publication Peer Review of the Biomedical Literature 2020.

38. Feltrin, A.S.; Tahira, A.C.; Simões, S.N.; Brentani, H.; Martins, D.C. Jr Assessment of complementarity of WGCNA and NERI results for identification of modules associated to schizophrenia spectrum disorders. PLoS One 2019, 14, e0210431. doi: 10.1371/journal.pone.0210431.

39. Guzzi, P.H.; Mercatelli, D.; Ceraolo, C.; Giorgi, F.M. Master Regulator Analysis of the SARS-CoV-2/Human Interactome. J. Clin. Med. Res. 2020, 9, doi: 10.3390/jcm9040982.

40. Garvin, M.R.; Alvarez, C.; Miller, J.I.; Prates, E.T.; Walker, A.M.; Amos, B.K.; Mast, A.E.; Justice, A.; Aronow, B.; Jacobson, D. A mechanistic model and therapeutic interventions for COVID-19 involving a RAS-mediated bradykinin storm. Elife 2020, 9, doi: 10.7554/eLife.59177.

41. Ehrenfeld, M.; Tincani, A.; Andreoli, L.; Cattalini, M.; Greenbaum, A.; Kanduc, D.; Alijotas-Reig, J.; Zinserling, V.; Semenova, N.; Amital, H.; et al. Covid-19 and autoimmunity. Autoimmunity Reviews 2020, 19, 102597.

42. Yan, J.; Risacher, S.L.; Shen, L.; Saykin, A.J. Network approaches to systems biology analysis of complex disease: integrative methods for multi-omics data. Brief. Bioinform. 2018, 19, 1370–1381, doi: 10.1093/bib/bbx066.

43. Kim, Y.-A.; Wuchty, S.; Przytycka, T.M. Identifying Causal Genes and Dysregulated Pathways in Complex Diseases. PLoS Computational Biology 2011, 7, e1001095.

44. Suratanee, A.; Plaimas, K. Network-based association analysis to infer new disease-gene relationships using large-scale protein interactions. PLOS ONE 2018, 13, e0199435.

45. Sadegh, S.; Matschinske, J.; Blumenthal, D.B.; Galindez, G.; Kacprowski, T.; List, M.; Nasirigerdeh, R.; Oubounyt, M.; Pichlmair, A.; Rose, T.D.; et al. Exploring the SARS-CoV-2 virus-host-drug interactome for drug repurposing. Nat. Commun. 2020, 11, 3518, doi: 10.1038/s41467-020-17189-2.

46. Maroli, N.; Bhasuran, B.; Natarajan, J.; Kolandaivel, P. The potential role of procyanidin as a therapeutic agent against SARS-CoV-2: a text mining, molecular docking and molecular dynamics simulation approach. J. Biomol. Struct. Dyn. 2020, 1–16, doi: 10.1080/07391102.2020.1823887.

47. Muthuramalingam, P.; Jeyasri, R.; Valliammai, A.; Selvaraj, A.; Karthika, C.; Gowrishankar, S.; Pandian, S.K.; Ramesh, M.; Chen, J.-T. Global multi-omics and systems pharmacological strategy unravel the multi-targeted therapeutic potential of natural bioactive molecules against COVID-19: An in silico approach. Genomics 2020, 112, 4486–4504.

48. Zheng, Y.-Y.; Ma, Y.-T.; Zhang, J.-Y.; Xie, X. COVID-19 and the cardiovascular system. Nat. Rev. Cardiol. 2020, 17, 259–260, doi: 10.1038/s41569-020-0360-5.

49. Liu, X.; Chen, Y.; Tang, W.; Zhang, L.; Chen, W.; Yan, Z.; Yuan, P.; Yang, M.; Kong, S.; Yan, L.; et al. Single-cell transcriptome analysis of the novel coronavirus (SARS-CoV-2) associated gene ACE2 expression in normal and non-obstructive azoospermia (NOA) human male testes. Sci. China Life Sci. 2020, 63, 1006–1015, doi: 10.1007/s11427-020-1705-0.

50. Ding, Y.; He, L.; Zhang, Q.; Huang, Z.; Che, X.; Hou, J.; Wang, H.; Shen, H.; Qiu, L.; Li, Z.; et al. Organ distribution of severe acute respiratory syndrome (SARS) associated coronavirus (SARS-CoV) in SARS patients: implications for pathogenesis and virus transmission pathways. J. Pathol. 2004, 203, 622–630, doi: 10.1002/path.1560.

51. Reis, F.M.; Bouissou, D.R.; Pereira, V.M.; Camargos, A.F.; dos Reis, A.M., Santos, R.A. Angiotensin-(1-7), its receptor Mas, and the angiotensin-converting enzyme type 2 are expressed in the human ovary. Fertility and Sterility 2011, 95, 176–181.

52. Somasundaram, N.P.; Ranathunga, I.; Ratnasamy, V.; Wijewickrama, P.S.A.; Dissanayake, H.A.; Yogendranathan, N.; Gamage, K.K.K.; de Silva, N.L.; Sumanatilleke, M.; Katulanda, P.; et al. The Impact of SARS-Cov-2 Virus Infection on the Endocrine System. J Endocr Soc 2020, 4, bvaa082. doi: 10.1210/jendso/bvaa082.

53. Ma, L.; Xie, W.; Li, D.; Shi, L.; Mao, Y.; Xiong, Y.; Zhang, Y.; Zhang, M. Effect of SARS-CoV-2 infection upon male gonadal function: A single center-based study. medRxiv 2020, 2020.03.21.20037267, doi: 10.1101/2020.03.21.20037267.

54. Fan, C.; Li, K.; Ding, Y.; Lu, W.L.; Wang, J. ACE2 Expression in Kidney and Testis May Cause Kidney and Testis Damage After 2019-nCoV Infection. Urology 2020.

55. Kaplan, S.A. Re: Urinary Frequency as a Possibly Overlooked Symptom in COVID-19 Patients: Does SARS-CoV-2 Cause Viral Cystitis? J. Urol. 2020, 101097JU000000000000124602, doi: 10.1097/JU.0000000000001246.02.

56. Sun, J.; Zhu, A.; Li, H.; Zheng, K.; Zhuang, Z.; Chen, Z.; Shi, Y.; Zhang, Z.; Chen, S.-B.; Liu, X.; et al. Isolation of infectious SARS-CoV-2 from urine of a COVID-19 patient. Emerg. Microbes Infect. 2020, 9, 991–993, doi: 10.1080/22221751.2020.1760144.

57. Mak, T.W.; Hauck, L.; Grothe, D.; Billia, F. p53 regulates the cardiac transcriptome. Proc. Natl. Acad. Sci. U. S. A. 2017, 114, 2331–2336, doi: 10.1073/pnas.1621436114.

58. Rhaleb, N.-E.; Yang, X.-P.; Carretero, O.A. The kallikrein-kinin system as a regulator of cardiovascular and renal function. Compr. Physiol. 2011, 1, 971–993, doi: 10.1002/cphy.c100053.

59. Isordia-Salas, I.; Pixley, R.A.; Sáinz, I.M.; Martínez-Murillo, C.; Colman, R.W. The role of plasma high molecular weight kininogen in experimental intestinal and systemic inflammation. Arch. Med. Res. 2004, 35, 369–377, doi: 10.1016/j.arcmed.2004.05.004.

60. Mattsson, E.; Herwald, H.; Cramer, H.; Persson, K.; Sjöbring, U.; Björck, L. Staphylococcus aureus induces release of bradykinin in human plasma. Infect. Immun. 2001, 69, 3877–3882, doi: 10.1128/IAI.69.6.3877-3882.2001.

61. Schreier, B.; Rabe, S.; Schneider, B.; Bretschneider, M.; Rupp, S.; Ruhs, S.; Neumann, J.; Rueckschloss, U.; Sibilia, M.; Gotthardt, M.; et al. Loss of epidermal growth factor receptor in vascular smooth muscle cells and cardiomyocytes causes arterial hypotension and cardiac hypertrophy. Hypertension 2013, 61, 333–340, doi: 10.1161/HYPERTENSIONAHA.112.196543.

62. Mitchell, H.D.; Eisfeld, A.J.; Stratton, K.G.; Heller, N.C.; Bramer, L.M.; Wen, J.; McDermott, J.E.; Gralinski, L.E.; Sims, A.C.; Le, M.Q.; et al. The Role of EGFR in Influenza Pathogenicity: Multiple Network-Based Approaches to Identify a Key Regulator of Non-lethal Infections. Front Cell Dev Biol 2019, 7, 200, doi: 10.3389/fcell.2019.00200.

63. Eimer, W.A.; Vijaya Kumar, D.K., Navalpur Shanmugam, N.K., Rodriguez, A.S.; Mitchell, T.; Washicosky, K.J.; György, B.; Breakefield, X.O.; Tanzi, R.E.; Moir, R.D. Alzheimer’s Disease-Associated β-Amyloid Is Rapidly Seeded by Herpesviridae to Protect against Brain Infection. Neuron 2018, 99, 56–63.e3, doi: 10.1016/j.neuron.2018.06.030.

64. Inyushin, M.; Zayas-Santiago, A.; Rojas, L.; Kucheryavykh, Y.; Kucheryavykh, L. Platelet-generated amyloid beta peptides in Alzheimer’s disease and glaucoma. Histol. Histopathol. 2019, 34, 843–856, doi: 10.14670/HH-18-111.

65. Kucheryavykh, L.Y.; Kucheryavykh, Y.V.; Washington, A.V.; Inyushin, M.Y. Amyloid Beta Peptide Is Released during Thrombosis in the Skin. Int. J. Mol. Sci. 2018, 19, doi: 10.3390/ijms19061705.

66. Chattopadhyay, S.; Basak, T.; Nayak, M.K.; Bhardwaj, G.; Mukherjee, A.; Bhowmick, R.; Sengupta, S.; Chakrabarti, O.; Chatterjee, N.S.; Chawla-Sarkar, M. Identification of cellular calcium binding protein calmodulin as a regulator of rotavirus A infection during comparative proteomic study. PLoS One 2013, 8, e56655, doi: 10.1371/journal.pone.0056655.

67. Lambert, D.W.; Clarke, N.E.; Hooper, N.M.; Turner, A.J. Calmodulin interacts with angiotensin-converting enzyme-2 (ACE2) and inhibits shedding of its ectodomain. FEBS Lett. 2008, 582, 385–390, doi: 10.1016/j.febslet.2007.11.085.

68. Montezano, A.C.; Cat, A.N.D.; Rios, F.J.; Touyz, R.M. Angiotensin II and Vascular Injury. Current Hypertension Reports 2014, 16.

69. Crackower, M.A.; Sarao, R.; Oudit, G.Y.; Yagil, C.; Kozieradzki, I.; Scanga, S.E.; Oliveira-dos-Santos, A.J.; da Costa, J.; Zhang, L.; Pei, Y.; et al. Angiotensin-converting enzyme 2 is an essential regulator of heart function. Nature 2002, 417, 822–828, doi: 10.1038/nature00786.

70. Zou, M.-X.; Liu, H.-Y.; Haraguchi, Y.; Soda, Y.; Tatemoto, K.; Hoshino, H. Apelin peptides block the entry of human immunodeficiency virus (HIV). FEBS Letters 2000, 473, 15–18.

71. Pagliaro, P.; Penna, C. ACE/ACE2 Ratio: A Key Also in 2019 Coronavirus Disease (Covid-19)? Front. Med. 2020, 7, 335, doi: 10.3389/fmed.2020.00335.

72. Wang, H.; Yang, P.; Liu, K.; Guo, F.; Zhang, Y.; Zhang, G.; Jiang, C. SARS coronavirus entry into host cells through a novel clathrin- and caveolae-independent endocytic pathway. Cell Res. 2008, 18, 290–301, doi: 10.1038/cr.2008.15.

73. Szczepanski, A.; Owczarek, K.; Milewska, A.; Baster, Z.; Rajfur, Z.; Mitchell, J.A.; Pyrc, K. Canine respiratory coronavirus employs caveolin-1-mediated pathway for internalization to HRT-18G cells. Vet. Res. 2018, 49, 55, doi: 10.1186/s13567-018-0551-9.

74. Emanuel, W.; Kirstin, M.; Vedran, F.; Asija, D.; Theresa, G.L.; Roberto, A.; Filippos, K.; David, K.; Salah, A.; Christopher, B.; et al. Bulk and single-cell gene expression profiling of SARS-CoV-2 infected human cell lines identifies molecular targets for therapeutic intervention.

75. Foy, B.H.; Carlson, J.C.T.; Reinertsen, E.; Padros I Valls, R., Pallares Lopez, R.; Palanques-Tost, E.; Mow, C.; Westover, M.B.; Aguirre, A.D.; Higgins, J.M. Association of Red Blood Cell Distribution Width With Mortality Risk in Hospitalized Adults With SARS-CoV-2 Infection. JAMA Netw Open 2020, 3, e2022058. doi: 10.1001/jamanetworkopen.2020.22058.

76. Qin, M.; Cao, Z.; Wen, J.; Yu, Q.; Liu, C.; Wang, F.; Zhang, J.; Yang, F.; Li, Y.; Fishbein, G.; et al. An Antioxidant Enzyme Therapeutic for COVID-19. Adv. Mater. 2020, e2004901, doi: 10.1002/adma.202004901.

77. Valiente-Alandi, I.; Potter, S.J.; Salvador, A.M.; Schafer, A.E.; Schips, T.; Carrillo-Salinas, F.; Gibson, A.M.; Nieman, M.L.; Perkins, C.; Sargent, M.A.; et al. Inhibiting Fibronectin Attenuates Fibrosis and Improves Cardiac Function in a Model of Heart Failure. Circulation 2018, 138, 1236–1252.

78. Torre, D.; Pugliese, A.; Ferrario, G.; Marietti, G.; Forno, B.; Zeroli, C. Interaction of human plasma fibronectin with viral proteins of human immunodeficiency virus. FEMS Immunol. Med. Microbiol. 1994, 8, 127–131, doi: 10.1111/j.1574-695X.1994.tb00434.x.

79. Barchetta, I.; Cavallo, M.G.; Baroni, M.G. COVID-19 and diabetes: Is this association driven by the DPP4 receptor? Potential clinical and therapeutic implications. Diabetes Research and Clinical Practice 2020, 163, 108165.

80. Li, K.; Wohlford-Lenane, C.L.; Channappanavar, R.; Park, J.-E.; Earnest, J.T.; Bair, T.B.; Bates, A.M.; Brogden, K.A.; Flaherty, H.A.; Gallagher, T.; et al. Mouse-adapted MERS coronavirus causes lethal lung disease in human DPP4 knockin mice. Proc. Natl. Acad. Sci. U. S. A. 2017, 114, E3119–E3128, doi: 10.1073/pnas.1619109114.

81. Fan, C.; Wu, X.; Liu, Q.; Li, Q.; Liu, S.; Lu, J.; Yang, Y.; Cao, Y.; Huang, W.; Liang, C.; et al. A Human DPP4-Knockin Mouse’s Susceptibility to Infection by Authentic and Pseudotyped MERS-CoV. Viruses 2018, 10, 448.

82. Iacobellis, G. COVID-19 and diabetes: Can DPP4 inhibition play a role? Diabetes Res. Clin. Pract. 2020, 162, 108125, doi: 10.1016/j.diabres.2020.108125.

83. Rose, C.E.; Edward Rose, C.; Sung, S.-S.J.; Fu, S.M. Significant Involvement of CCL2 (MCP-1) in Inflammatory Disorders of the Lung. Microcirculation 2003, 10, 273–288.

84. He, L.; Ding, Y.; Zhang, Q.; Che, X.; He, Y.; Shen, H.; Wang, H.; Li, Z.; Zhao, L.; Geng, J.; et al. Expression of elevated levels of pro-inflammatory cytokines in SARS-CoV-infected ACE2+ cells in SARS patients: relation to the acute lung injury and pathogenesis of SARS. J. Pathol. 2006, 210, 288–297, doi: 10.1002/path.2067.

85. Huang, C.; Wang, Y.; Li, X.; Ren, L.; Zhao, J.; Hu, Y.; Zhang, L.; Fan, G.; Xu, J.; Gu, X.; et al. Clinical features of patients infected with 2019 novel coronavirus in Wuhan, China. The Lancet 2020, 395, 497–506.

86. Zhang, L.; Sun, X.; Chen, S.; Yang, C.; Shi, B.; Zhou, L.; Zhao, J. Long noncoding RNA DANCR regulates -Smad 4 axis to promote chondrogenic differentiation of human synovium-derived mesenchymal stem cells. Biosci. Rep. 2017, 37, doi: 10.1042/BSR20170347.

87. Su, Y.; Feng, W.; Shi, J.; Chen, L.; Huang, J.; Lin, T. circRIP2 accelerates bladder cancer progression via miR-1305/Tgf-β2/smad3 pathway. Molecular Cancer 2020, 19.

88. Yang, X.; Letterio, J.J.; Lechleider, R.J.; Chen, L.; Hayman, R.; Gu, H.; Roberts, A.B.; Deng, C. Targeted disruption of SMAD3 results in impaired mucosal immunity and diminished T cell responsiveness to TGF-beta. EMBO J. 1999, 18, 1280–1291, doi: 10.1093/emboj/18.5.1280.

89. Mirzaei, H.; Faghihloo, E. Viruses as key modulators of the TGF-β pathway; a double-edged sword involved in cancer. Reviews in Medical Virology 2018, 28, e1967.

90. Eulertaimor, G.; Heger, J. The complex pattern of SMAD signalling in the cardiovascular system☆. Cardiovascular Research 2006, 69, 15–25.

91. Qiu, H.; Zhang, Y.; Zhao, Q.; Jiang, H.; Yan, J.; Liu, Y. Platelet miR-587 may be Used as a Potential Biomarker for Diagnosis of Patients with Acute Coronary Syndrome. Clinical Laboratory 2020, 66.

92. Tang, C.-M.; Zhang, M.; Huang, L.; Hu, Z.-Q.; Zhu, J.-N.; Xiao, Z.; Zhang, Z.; Lin, Q.-X.; Zheng, X.-L.; -Yang, M.; et al. CircRNA_000203 enhances the expression of fibrosis-associated genes by derepressing targets of miR-26b-5p, Col1a2 and CTGF, in cardiac fibroblasts. Scientific Reports 2017, 7.

93. Chen, X.; Zhou, L.; Peng, N.; Yu, H.; Li, M.; Cao, Z.; Lin, Y.; Wang, X.; Li, Q.; Wang, J.; et al. MicroRNA-302a suppresses influenza A virus–stimulated interferon regulatory factor-5 expression and cytokine storm induction. Journal of Biological Chemistry 2017, 292, 21291–21303.

94. Hamada-Tsutsumi, S.; Naito, Y.; Sato, S.; Takaoka, A.; Kawashima, K.; Isogawa, M.; Ochiya, T.; Tanaka, Y. The antiviral effects of human microRNA miR-302c-3p against hepatitis B virus infection. Alimentary Pharmacology & Therapeutics 2019, 49, 1060–1070.

95. Peng, N.; Yang, X.; Zhu, C.; Zhou, L.; Yu, H.; Li, M.; Lin, Y.; Wang, X.; Li, Q.; She, Y.; et al. MicroRNA-302 Cluster Downregulates Enterovirus 71–Induced Innate Immune Response by Targeting KPNA2. The Journal of Immunology 2018, 201, 145–156.

96. Li, G.; Song, Y.; Li, Y.-D.; Jie, L.-J.; Wu, W.-Y.; Li, J.-Z.; Zhang, Q.; Wang, Y. Circulating miRNA-302 family members as potential biomarkers for the diagnosis of acute heart failure. Biomarkers in Medicine 2018, 12, 871–880.

97. Braga, L.; Ali, H.; Secco, I.; Giacca, M. Non-coding RNA therapeutics for cardiac regeneration. Cardiovasc. Res. 2020, doi: 10.1093/cvr/cvaa071.

98. Wang, J.; Huang, R.; Xu, Q.; Zheng, G.; Qiu, G.; Ge, M.; Shu, Q.; Xu, J. Mesenchymal Stem Cell–Derived Extracellular Vesicles Alleviate Acute Lung Injury Via Transfer of miR-27a-3p. Critical Care Medicine 2020, 1.

99. Zhao, X.-R.; Zhang, Z.; Gao, M.; Li, L.; Sun, P.-Y.; Xu, L.-N.; Qi, Y.; Yin, L.-H.; Peng, J.-Y. MicroRNA-27a-3p aggravates renal ischemia/reperfusion injury by promoting oxidative stress via targeting growth factor receptor-bound protein 2. Pharmacol. Res. 2020, 155, 104718, doi: 10.1016/j.phrs.2020.104718.

100. Lv, X.; Yan, J.; Jiang, J.; Zhou, X.; Lu, Y.; Jiang, H. MicroRNA-27a-3p suppression of peroxisome proliferator-activated receptor-γ contributes to cognitive impairments resulting from sevoflurane treatment. Journal of Neurochemistry 2017, 143, 306–319.

101. Vegter, E.L.; Ovchinnikova, E.S.; van Veldhuisen, D.J.; Jaarsma, T.; Berezikov, E.; van der Meer, P.; Voors, A.A. Low circulating microRNA levels in heart failure patients are associated with atherosclerotic disease and cardiovascular-related rehospitalizations. Clinical Research in Cardiology 2017, 106, 598–609.

102. Ovchinnikova, E.S.; Schmitter, D.; Vegter, E.L.; ter Maaten, J.M., Valente, M.A.E.; Liu, L.C.Y.; van der Harst, P.; Pinto, Y.M.; de Boer, R.A.; Meyer, S.; et al. Signature of circulating microRNAs in patients with acute heart failure. European Journal of Heart Failure 2016, 18, 414–423.

103. Yamada, K.; Takizawa, S.; Ohgaku, Y.; Asami, T.; Furuya, K.; Yamamoto, K.; Takahashi, F.; Hamajima, C.; Inaba, C.; Endo, K.; et al. MicroRNA 16-5p is upregulated in calorie-restricted mice and modulates inflammatory cytokines of macrophages. Gene 2020, 725, 144191, doi: 10.1016/j.gene.2019.144191.

104. Ye, E.-A.; Liu, L.; Jiang, Y.; Jan, J.; Gaddipati, S.; Suvas, S.; Steinle, J.J. miR-15a/16 reduces retinal leukostasis through decreased pro-inflammatory signalling. J. Neuroinflammation 2016, 13, 305, doi: 10.1186/s12974-016-0771-8.

105. Mariconti, M.; Vola, A.; Manciulli, T.; Genco, F.; Lissandrin, R.; Meroni, V.; Rosenzvit, M.; Tamarozzi, F.; Brunetti, E. Correction to: Role of microRNAs in host defense against Echinococcus granulosus infection: a preliminary assessment. Immunol. Res. 2019, 67, 98, doi: 10.1007/s12026-018-9060-1.

106. Biswas, S.; Haleyurgirisetty, M.; Lee, S.; Hewlett, I.; Devadas, K. Development and validation of plasma miRNA biomarker signature panel for the detection of early HIV-1 infection. EBioMedicine 2019, 43, 307–316, doi: 10.1016/j.ebiom.2019.04.023.

107. Ketprasit, N.; Cheng, I.S.; Deutsch, F.; Tran, N.; Imwong, M.; Combes, V.; Palasuwan, D. The characterization of extracellular vesicles-derived microRNAs in Thai malaria patients. Malar. J. 2020, 19, 285, doi: 10.1186/s12936-020-03360-z.

108. Jia, H.-L.; He, C.-H.; Wang, Z.-Y.; Xu, Y.-F.; Yin, G.-Q.; Mao, L.-J.; Liu, C.-W.; Deng, L. MicroRNA expression profile in exosome discriminates extremely severe infections from mild infections for hand, foot and mouth disease. BMC Infect. Dis. 2014, 14, 506, doi: 10.1186/1471-2334-14-506.

109. Marquez-Pedroza, J.; Cárdenas-Bedoya, J.; Morán-Moguel, M.C.; Escoto-Delgadillo, M.; Torres-Mendoza, B.M.; Pérez-Ríos, A.M.; González-Enriquez, G.V.; Vázquez-Valls, E. Plasma microRNA expression levels in HIV-1-positive patients receiving antiretroviral therapy. Biosci. Rep. 2020, 40, doi: 10.1042/BSR20194433.

110. Zhao, Y.; Yan, M.; Chen, C.; Gong, W.; Yin, Z.; Li, H.; Fan, J.; Zhang, X.A.; Wang, D.W.; Zuo, H. MiR-124 aggravates failing hearts by suppressing CD151-facilitated angiogenesis in heart. Oncotarget 2018, 9, 14382–14396, doi: 10.18632/oncotarget.24205.

111. Dang, J.W.; Tiwari, S.K.; Qin, Y.; Rana, T.M. Genome-wide Integrative Analysis of Zika-Virus-Infected Neuronal Stem Cells Reveals Roles for MicroRNAs in Cell Cycle and Stemness. Cell Rep. 2019, 27, 3618–3628.e5, doi: 10.1016/j.celrep.2019.05.059.

