## Supplementary Table 2 for "ACE2 interaction networks in COVID-19: a physiological framework for prediction of outcome in patients with cardiovascular risk factors"

**Supplementary Table 2. Description of the genes from the ACE2 interaction network**. Fields Tissue: Heart, Tissue: Lungs and Tissue: Nervous System are associated with gene expression confidence from TISSUES database <https://tissues.jensenlab.org/> . Columns marked as GO (gene Ontology) show association of the genes with selected biological processes related to heart failure.

Complete ACE network in interactive form is available here: http://www.ndexbio.org/#/network/c5c336c2-9ab3-11ea-aaef-0ac135e8bacf

| **Gene Symbol** | **Degree of connections** | **Tissue: Heart** | **Tissue: Lungs** | **Tissue: Nervous System** | **GO: Atherosclerosis** | **GO: Cardiovascular disease** | **GO: Diabetes** | **GO: Dyslipidemia** | **GO: Nephophathy** | **GO: Neuropathy** | **GO: Virus-related protein** | **Description** |
| --- | --- | --- | --- | --- | --- | --- | --- | --- | --- | --- | --- | --- |
| ACE2 | 49 | 4.63 | 4.47 | 4.47 | **-** | **Y** | **-** | **-** | **Y** | **-** | **Y** | **Description:** Angiotensin-converting enzyme homolog; Carboxypeptidase which converts angiotensin I to angiotensin 1-9, a peptide of unknown function, and angiotensin II to angiotensin 1-7, a vasodilator. Also able to hydrolyze apelin- 13 and dynorphin-13 with high efficiency.  **COVID-19 related function:** May be an important regulator of heart function. ACE2 is highly expressed in ventricular cardiomyocytes, heart macrophages, pericytes of the heart, lung, and kidney. ACE2 is able to modify acute and chronic inflammatory response and maintain ROS generation by several pathways, such as Ang-(1-7)/Mas, NF-κB, MAPK, AT1/ERK1/2, TGF-β/SMAD, JNK1/2, PI3K-Akt/ eNOS, ANGII/NADPH, PKC/p38 MAPK, MMP2, MMP9. Abnormalities of ACE2 pathway signaling are closely related to pathophysiology of hypertension, myocardial tissue fibrosis, inflammation and therefore leads to cardiac remodeling and heart failure. Moreover, dysregulation of ACE2 function mediates renal damage and fibrosis and is markedly related to pulmonary inflammation, lung injury and severe acute lung failure.  **Ref:** [1–13];  **Gene Ontology:** cardiac muscle functions; inflammation |
| ACE | 33 | 4.74 | 4.62 | 4.90 | **-** | **Y** | **Y** | **-** | **Y** | **Y** | **-** | **Description:** Angiotensin I converting enzyme; Converts angiotensin I to angiotensin II by release of the terminal His-Leu, this results in an increase of the vasoconstrictor activity of angiotensin. Also able to inactivate bradykinin, a potent vasodilator. Has also a glycosidase activity which releases GPI-anchored proteins from the membrane by cleaving the mannose linkage in the GPI moiety; CD molecules;  **Gene Ontology:** |
| REN | 32 | 3.47 | 2.24 | 3.21 | **Y** | **Y** | **Y** | **-** | **-** | **-** | **-** | **Description:** Angiotensinogenase; Renin is a highly specific endopeptidase, whose only known function is to generate angiotensin I from angiotensinogen in the plasma, initiating a cascade of reactions that produce an elevation of blood pressure and increased sodium retention by the kidney.  **Gene Ontology:** |
| INS | 31 | 3.40 | 2.53 | 3.47 | **Y** | **Y** | **Y** | **-** | **-** | **-** | **-** | **Description:** Insulin; Insulin decreases blood glucose concentration. It increases cell permeability to monosaccharides, amino acids and fatty acids. It accelerates glycolysis, the pentose phosphate cycle, and glycogen synthesis in liver.  **COVID-19 related function:** Dysregulation of insulin signaling results in oxidative stress formation and subsequently to blunted calcium handling by sarcoplasmic reticulum as well as impaired mitochondrial, endothelial and cardiac dysfunction. Consequently, these abnormalities result in cardiac hypertrophy, fibrosis, cardiomyocyte apoptosis and heart failure. Hyperinsulinemia is associated with pro-hypertensive effects of inusline, leading to RAS/MAP mediated vascular smooth muscle proliferation and exacerbated atherosclerosis. Impaired insulin action synergistically with hypertension leads to cardiovascular complications. On the other hand, overactivity of RAS may dysregulate insulin response via Ang II/P13-kinase mediated pathway contributing to insulin resistance. . **Ref:** [71–75]  **Gene Ontology:** |
| KNG1 | 30 | 3.77 | 2.77 | 4.64 | **-** | **-** | **-** | **-** | **-** | **-** | **-** | **Description:** Williams-Fitzgerald-Flaujeac factor; (1) Kininogens are inhibitors of thiol proteases; (2) HMW-kininogen plays an important role in blood coagulation by helping to position optimally prekallikrein and factor XI next to factor XII; (3) HMW-kininogen inhibits the thrombin- and plasmin- induced aggregation of thrombocytes; (4) the active peptide bradykinin that is released from HMW-kininogen shows a variety of physiological effects: (4A) influence in smooth muscle contraction, (4B) induction of hypotension, (4C) natriuresis and diuresis, (4D) decrease in blood glucose level, (4E) it is a mediator of inflammation and causes (4E1) increase in vascular permeability, (4E2) stimulation of nociceptors (4E3) release of other mediators of inflammation (e.g. prostaglandins), (4F) it has a cardioprotective effect (directly via bradykinin action, indirectly via endothelium-derived relaxing factor action); (5) LMW-kininogen inhibits the aggregation of thrombocytes; (6) LMW- kininogen is in contrast to HMW-kininogen not involved in blood clotting; Cystatins, type 3;  **Gene Ontology:** coagulation; inflammation |
| AGT | 28 | 4.65 | 2.58 | 5.00 | **-** | **-** | **-** | **-** | **Y** | **Y** | **-** | **Description:** Angiotensinogen (serpin peptidase inhibitor, clade A, member 8); Essential component of the renin-angiotensin system (RAS), a potent regulator of blood pressure, body fluid and electrolyte homeostasis; Endogenous ligands;  **Gene Ontology:** angiogenesis; cardiac muscle functions; fibrosis; inflammation; muscle hypertrophy |
| AGTR1 | 26 | 3.07 | 1.94 | 4.55 | **-** | **-** | **-** | **-** | **Y** | **-** | **-** | **Description:** Angiotensin II receptor, type 1; Receptor for angiotensin II. Mediates its action by association with G proteins that activate a phosphatidylinositol- calcium second messenger system.  **Gene Ontology:** angiogenesis; inflammation |
| MME | 24 | 2.79 | 2.86 | 4.61 | **-** | **-** | **-** | **-** | **-** | **Y** | **-** | **Description:** Common acute lymphocytic leukemia antigen; Thermolysin-like specificity, but is almost confined on acting on polypeptides of up to 30 amino acids. Biologically important in the destruction of opioid peptides such as Met- and Leu-enkephalins by cleavage of a Gly-Phe bond. Able to cleave angiotensin-1, angiotensin-2 and angiotensin 1-9. Involved in the degradation of atrial natriuretic factor (ANF). Displays UV-inducible elastase activity toward skin preelastic and elastic fibers; Belongs to the peptidase M13 family.  **Gene Ontology:** |
| NOS3 | 22 | 3.67 | 2.53 | 4.62 | **Y** | **Y** | **Y** | **Y** | **Y** | **Y** | **-** | **Description:** Nitric oxide synthase 3 (endothelial cell); Produces nitric oxide (NO) which is implicated in vascular smooth muscle relaxation through a cGMP-mediated signal transduction pathway. NO mediates vascular endothelial growth factor (VEGF)-induced angiogenesis in coronary vessels and promotes blood clotting through the activation of platelets.  **Gene Ontology:** angiogenesis |
| DPP4 | 22 | 2.89 | 2.59 | 2.64 | **Y** | **Y** | **Y** | **Y** | **Y** | **Y** | **Y** | **Description:** Adenosine deaminase complexing protein 2; Cell surface glycoprotein receptor involved in the costimulatory signal essential for T-cell receptor (TCR)-mediated T-cell activation. Acts as a positive regulator of T-cell coactivation, by binding at least ADA, CAV1, IGF2R, and PTPRC. Its binding to CAV1 and CARD11 induces T-cell proliferation and NF- kappa-B activation in a T-cell receptor/CD3-dependent manner. Its interaction with ADA also regulates lymphocyte-epithelial cell adhesion. In association with FAP is involved in the pericellular proteolysis of the extracellular matrix (ECM), the migration and invasion of endothelial cells into the ECM. May be involved in the promotion of lymphatic endothelial cells adhesion, migration and tube formation. When overexpressed, enhanced cell proliferation, a process inhibited by GPC3. Acts also as a serine exopeptidase with a dipeptidyl peptidase activity that regulates various physiological processes by cleaving peptides in the circulation, including many chemokines, mitogenic growth factors, neuropeptides and peptide hormones. Removes N-terminal dipeptides sequentially from polypeptides having unsubstituted N-termini provided that the penultimate residue is proline; Belongs to the peptidase S9B family. DPPIV subfamily.  **COVID-19 related function:** High expression of DPP4 can be found among patients with obesity, metabolic syndrome and T2DM, who are at risk of a severe course of COVID-19. The DPP4 gene encodes dipeptidyl peptidase 4, which is involved in maintaining glucose and insulin metabolism in diabetes, as well as in immune regulation and inflammatory response via TGF-β1, MAPK, NF-κB pathways. DPP4 mediates costimulatory signal essential for T-cell receptor-mediated T-cell activation and proliferation. DPP4 mediates pro-inflammatory cytokine production in a model of lung injury. Moreover, DPP4 is also involved in atherosclerosis and maintaining cardiovascular homeostasis, endothelial repair, inflammatory response and ischemic injury of vascular smooth muscle. DPP4 was found as a functional receptor for Middle East respiratory syndrome coronavirus (MERS-CoV). DPP4 may be involved in anosomia which occurs commonly among COVID-19 patients. **Ref:** [14–22];  **Gene Ontology:** |
| ENPEP | 20 | 2.46 | 2.07 | 2.72 | **-** | **-** | **-** | **-** | **-** | **-** | **-** | **Description:** Glutamyl aminopeptidase (aminopeptidase A)/ Differentiation Antigen Gp160; Appears to have a role in the catabolic pathway of the renin-angiotensin system. Probably plays a role in regulating growth and differentiation of early B-lineage cells; Belongs to the peptidase M1 family.  **Gene Ontology:** angiogenesis |
| ANPEP | 19 | 2.73 | 2.54 | 2.49 | **-** | **-** | **-** | **-** | **-** | **-** | **Y** | **Description:** Alanyl Aminopeptidase, Membrane / Myeloid plasma membrane glycoprotein CD13; Broad specificity aminopeptidase which plays a role in the final digestion of peptides generated from hydrolysis of proteins by gastric and pancreatic proteases. Also involved in the processing of various peptides including peptide hormones, such as angiotensin III and IV, neuropeptides, and chemokines. May also be involved the cleavage of peptides bound to major histocompatibility complex class II molecules of antigen presenting cells. May have a role in angiogenesis and promote cholesterol crystallization; Aminopeptidases;  **COVID-19 related function:** Aminopeptidase N plays a role in the processing of various peptides including peptide hormones, such as angiotensin III and IV, neuropeptides, and chemokines. APN serve as a receptor for the HCoV-229E as well as other non-human coronaviruses. APN may influence blood pressure by regulating the metabolism of Ang III. APN is associated with inflammatory response, angiogenesis and tumor growth and is related with several autoimmune diseases, such as rheumatoid arthritis, psoriasis and inflammatory bowel disease and various types of leukemia and lymphoma.**Ref:** [23–28];  **Gene Ontology:** angiogenesis |
| EDN1 | 18 | 3.42 | 3.08 | 3.74 | **Y** | **-** | **-** | **Y** | **Y** | **Y** | **-** | **Description:** Preproendothelin-1; Endothelins are endothelium-derived vasoconstrictor peptides; Belongs to the endothelin/sarafotoxin family.  **Gene Ontology:** cardiac muscle functions; coagulation; muscle hypertrophy |
| NTS | 17 | 1.93 | 1.90 | 4.77 | **-** | **Y** | **-** | **-** | **-** | **-** | **-** | **Description:** Neurotensin/neuromedin N; Neurotensin may play an endocrine or paracrine role in the regulation of fat metabolism. It causes contraction of smooth muscle; Belongs to the neurotensin family.  **Gene Ontology:** |
| AGTR2 | 16 | 2.50 | 4.49 | 2.59 | **-** | **-** | **-** | **-** | **-** | **-** | **-** | **Description:** Angiotensin II receptor, type 2; Receptor for angiotensin II. Cooperates with MTUS1 to inhibit ERK2 activation and cell proliferation.  **Gene Ontology:** cardiac muscle functions; fibrosis; inflammation |
| THOP1 | 15 | 2.15 | 4.40 | 4.74 | **-** | **-** | **-** | **-** | **-** | **-** | **-** | **Description:** Thimet oligopeptidase 1; Involved in the metabolism of neuropeptides under 20 amino acid residues long. Involved in cytoplasmic peptide degradation. Able to degrade the amyloid-beta precursor protein and generate amyloidogenic fragments; M3 metallopeptidases;  **Gene Ontology:** |
| CCL2 | 15 | 3.73 | 3.51 | 3.85 | **Y** | **-** | **Y** | **-** | **Y** | **Y** | **Y** | **Description:** Monocyte chemotactic and activating factor; Chemotactic factor that attracts monocytes and basophils but not neutrophils or eosinophils. Augments monocyte anti-tumor activity. Has been implicated in the pathogenesis of diseases characterized by monocytic infiltrates, like psoriasis, rheumatoid arthritis or atherosclerosis. May be involved in the recruitment of monocytes into the arterial wall during the disease process of atherosclerosis; Belongs to the intercrine beta (chemokine CC) family.  **COVID-19 related function:** CCL2 is involved in immunoregulatory and inflammatory processes maintaining chemotactic activity for monocytes and basophils. CCL2 has been found to maintain inflammatory response and cytokine storm in SARS patients. CCL2 expression levels are constantly elevated among severe SARS patients in comparison to non-severe controls and associated with diffuse pulmonary infiltrates, higher body temperatures and longer hospitalization. CCL2 is also involved in pathogenesis of HIV-1, diabetes, ischemia-related neuronal death and cardiovascular diseases, especially coronary artery disease and atherosclerosis. CCL2 protein has been implicated in lung inflammatory disorders and contributes to development of pulmonary fibrosis.**Ref:** [29–36];  **Gene Ontology:** angiogenesis; fibrosis; inflammation |
| XPNPEP2 | 14 | 1.36 | 4.14 | 1.47 | **-** | **-** | **-** | **-** | **-** | **Y** | **-** | **Description:** X-prolyl aminopeptidase (aminopeptidase P) 2, membrane-bound; Membrane-bound metalloprotease which catalyzes the removal of a penultimate prolyl residue from the N-termini of peptides, such as Arg-Pro-Pro. May play a role in the metabolism of the vasodilator bradykinin; Belongs to the peptidase M24B family.  **Gene Ontology:** |
| PRCP | 14 | 2.79 | 4.36 | 4.83 | **Y** | **Y** | **-** | **-** | **Y** | **-** | **-** | **Description:** Prolylcarboxypeptidase (angiotensinase C); Cleaves C-terminal amino acids linked to proline in peptides such as angiotensin II, III and des-Arg9-bradykinin. This cleavage occurs at acidic pH, but enzymatic activity is retained with some substrates at neutral pH; M14 carboxypeptidases;  **Gene Ontology:** angiogenesis; coagulation |
| PREP | 14 | 3.15 | 1.92 | 4.00 | **-** | **-** | **-** | **-** | **-** | **-** | **-** | **Description:** Post-proline cleaving enzyme; Cleaves peptide bonds on the C-terminal side of prolyl residues within peptides that are up to approximately 30 amino acids long; Belongs to the peptidase S9A family.  **Gene Ontology:** |
| CMA1 | 13 | 4.57 | 2.44 | 2.18 | **-** | **-** | **-** | **-** | **Y** | **Y** | **-** | **Description:** Chymase 1, mast cell; Major secreted protease of mast cells with suspected roles in vasoactive peptide generation, extracellular matrix degradation, and regulation of gland secretion.  **Gene Ontology:** angiogenesis; inflammation |
| APLN | 13 | 2.29 | 0.96 | 4.40 | **-** | **-** | **Y** | **-** | **-** | **Y** | **-** | **Description:** Apelin; Endogenous ligands;  **Gene Ontology:** angiogenesis; fibrosis |
| CAT | 13 | 3.74 | 3.41 | 4.84 | **-** | **-** | **-** | **-** | **Y** | **-** | **-** | **Description:** Catalase; Occurs in almost all aerobically respiring organisms and serves to protect cells from the toxic effects of hydrogen peroxide. Promotes growth of cells including T-cells, B-cells, myeloid leukemia cells, melanoma cells, mastocytoma cells and normal and transformed fibroblast cells; Belongs to the catalase family.  **Gene Ontology:** |
| TGFB1 | 11 | 3.20 | 3.19 | 3.39 | **-** | **Y** | **-** | **-** | **Y** | **-** | **-** | **Description:** Transforming growth factor, beta 1; Multifunctional protein that controls proliferation, differentiation and other functions in many cell types. Many cells synthesize TGFB1 and have specific receptors for it. It positively and negatively regulates many other growth factors. It plays an important role in bone remodeling as it is a potent stimulator of osteoblastic bone formation, causing chemotaxis, proliferation and differentiation in committed osteoblasts (By similarity). Stimulates sustained production of collagen through the activation of CREB3L1 by regulated intramembrane proteolysis (RIP). Can promote either T-helper 17 cells (Th17) or regulatory T-cells (Treg) lineage differentiation in a concentration-dependent manner. At high concentrations, leads to FOXP3-mediated suppression of RORC and down-regulation of IL-17 expression, favoring Treg cell development. At low concentrations in concert with IL-6 and IL-21, leads to expression of the IL-17 and IL-23 receptors, favoring differentiation to Th17 cells. Mediates SMAD2/3 activation by inducing its phosphorylation and subsequent translocation to the nucleus. Can induce epithelial-to-mesenchymal transition (EMT) and cell migration in various cell types; Endogenous ligands;  **Gene Ontology:** cardiac muscle functions; fibrosis; inflammation |
| TFRC | 11 | 3.32 | 3.32 | 4.94 | **-** | **-** | **-** | **-** | **-** | **-** | **Y** | **Description:** Transferrin receptor protein 1; Cellular uptake of iron occurs via receptor-mediated endocytosis of ligand-occupied transferrin receptor into specialized endosomes. Endosomal acidification leads to iron release. The apotransferrin-receptor complex is then recycled to the cell surface with a return to neutral pH and the concomitant loss of affinity of apotransferrin for its receptor. Transferrin receptor is necessary for development of erythrocytes and the nervous system (By similarity). A second ligand, the heditary hemochromatosis protein HFE, competes for binding with transferrin for an overlapping C-terminal binding site. Positively regulates T and B cell proliferation through iron uptake; Belongs to the peptidase M28 family. M28B subfamily.  **COVID-19 related function:** Transferrin receptor necessary for cellular iron uptake essential in erythropoiesis, neurologic development and adaptive immunity. Positively regulates T and B cell function. Mutation in TFRC causes combined immunodeficiency an is involved in pathogenesis of anemia.**Ref:** [37];  **Gene Ontology:** |
| APLNR | 10 | 2.83 | 1.80 | 4.95 | **-** | **-** | **-** | **-** | **Y** | **Y** | **-** | **Description:** G-protein coupled receptor HG11; Receptor for apelin receptor early endogenous ligand (APELA) and apelin (APLN) hormones coupled to G proteins that inhibit adenylate cyclase activity. Plays a key role in early development such as gastrulation and heart morphogenesis by acting as a receptor for APELA hormone (By similarity). Plays also a role in various processes in adults such as regulation of blood vessel formation, blood pressure, heart contractility, and heart failure by acting as a receptor for APLN hormone; Belongs to the G-protein coupled receptor 1 family.  **Gene Ontology:** angiogenesis |
| ATP6AP2 | 9 | 2.88 | 3.27 | 4.99 | **-** | **-** | **-** | **-** | **-** | **Y** | **-** | **Description:** Vacuolar ATP synthase membrane sector-associated protein M8-9; Functions as a renin and prorenin cellular receptor. May mediate renin-dependent cellular responses by activating ERK1 and ERK2. By increasing the catalytic efficiency of renin in AGT/angiotensinogen conversion to angiotensin I, it may also play a role in the renin-angiotensin system (RAS).  **Gene Ontology:** |
| MEP1A | 9 | 1.27 | 0.77 | 2.03 | **-** | **-** | **-** | **-** | **-** | **-** | **Y** | **Description:** N-benzoyl-L-tyrosyl-P-amino-benzoic acid hydrolase subunit alpha; Meprin A subunit alpha; Belongs to the peptidase M12A family.  **COVID-19 related function:** MEP1A encodes The metalloproteases meprin A which is involved in cell proliferation, angiogenesis, carcinogenesis. Meprin A is involved in inflammatory response and pathogenesis of IBD, inflammatory lung disease, and renal injury. Gene variants of MEP1A might be associated with glucose metabolism and insulin sensitivity. Additionally, meprin A may interact with the RAS axis. Increased expression of meprins is associated with fibrotic conditions in the skin and the lung fibrosis, pulmonary hypertension, Alzheimer disease and protects against sepsis.**Ref:** [38–41]; |
| MEP1B | 9 | 0.93 | 0.51 | 0.99 | **-** | **-** | **-** | **-** | **Y** | **-** | **-** | **Description:** N-benzoyl-L-tyrosyl-P-amino-benzoic acid hydrolase subunit beta; Membrane metallopeptidase that sheds many membrane-bound proteins. Exhibits a strong preference for acidic amino acids at the P1' position. Known substrates include: FGF19, VGFA, IL1B, IL18, procollagen I and III, E-cadherin, KLK7, gastrin, ADAM10, tenascin-C. The presence of several pro-inflammatory cytokine among substrates implicate MEP1B in inflammation. It is also involved in tissue remodeling due to its capability to degrade extracellular matrix components; Astacins;  **Gene Ontology:** inflammation |
| MAS1 | 9 | 1.66 | 1.63 | 2.31 | **-** | **-** | **-** | **-** | **-** | **-** | **-** | **Description:** MAS1 proto-oncogene, G protein-coupled receptor; Receptor for angiotensin 1-7 (By similarity). Acts specifically as a functional antagonist of AGTR1 (angiotensin-2 type 1 receptor), although it up-regulates AGTR1 receptor levels. Positive regulation of AGTR1 levels occurs through activation of the G-proteins GNA11 and GNAQ, and stimulation of the protein kinase C signaling cascade. The antagonist effect on AGTR1 function is probably due to AGTR1 being physically altered by MAS1.  **Gene Ontology:** inflammation |
| CTGF | 9 | 4.74 | 4.21 | 4.61 | **-** | **-** | **-** | **-** | **-** | **Y** | **-** | **Description:** Hypertrophic chondrocyte-specific protein 24; Major connective tissue mitoattractant secreted by vascular endothelial cells. Promotes proliferation and differentiation of chondrocytes. Mediates heparin- and divalent cation-dependent cell adhesion in many cell types including fibroblasts, myofibroblasts, endothelial and epithelial cells. Enhances fibroblast growth factor-induced DNA synthesis; Belongs to the CCN family.  **Gene Ontology:** angiogenesis; cardiac muscle functions; fibrosis |
| CTSG | 8 | 2.69 | 2.53 | 2.17 | **-** | **-** | **-** | **-** | **-** | **-** | **-** | **Description:** Cathepsin G; Serine protease with trypsin- and chymotrypsin-like specificity. Cleaves complement C3. Has antibacterial activity against the Gram-negative bacterium P.aeruginosa, antibacterial activity is inhibited by LPS from P.aeruginosa, Z-Gly-Leu-Phe- CH2Cl and phenylmethylsulfonyl fluoride; Belongs to the peptidase S1 family.  **Gene Ontology:** |
| NLN | 8 | 1.93 | 1.19 | 4.71 | **-** | **-** | **-** | **-** | **-** | **-** | **-** | **Description:** Neurolysin (metallopeptidase M3 family); Hydrolyzes oligopeptides such as neurotensin, bradykinin and dynorphin A; Belongs to the peptidase M3 family.  **Gene Ontology:** |
| SLC15A1 | 7 | 1.28 | 1.65 | 2.01 | **-** | **-** | **-** | **-** | **-** | **-** | **-** | **Description:** Solute carrier family 15 (oligopeptide transporter), member 1; Proton-coupled intake of oligopeptides of 2 to 4 amino acids with a preference for dipeptides. May constitute a major route for the absorption of protein digestion end-products; Solute carriers;  **Gene Ontology:** |
| ADAM17 | 7 | 2.81 | 2.67 | 2.83 | **Y** | **-** | **-** | **-** | **Y** | **Y** | **Y** | **Description:** Disintegrin and metalloproteinase domain-containing protein 17; Cleaves the membrane-bound precursor of TNF-alpha to its mature soluble form. Responsible for the proteolytical release of soluble JAM3 from endothelial cells surface. Responsible for the proteolytic release of several other cell-surface proteins, including p75 TNF-receptor, interleukin 1 receptor type II, p55 TNF-receptor, transforming growth factor-alpha, L-selectin, growth hormone receptor, MUC1 and the amyloid precursor protein. Acts as an activator of Notch pathway by mediating cleavage of Notch, generating the membrane-associated intermediate fragment called Notch extracellular truncation (NEXT). Plays a role in the proteolytic processing of ACE2; ADAM metallopeptidase domain containing;  **COVID-19 related function:** ADAM 17 is metalloprotease involved in cell-cell and cell-matrix interactions, including fertilization, muscle development, and neurogenesis. Adam 17 mediates inflammatory response by cleaving cytokines, cytokine receptors ligands of ErbB and adhesion proteins therefore play a role in chemotaxis, fever induction and T-cell activation. On the other hand Adam 17 also mediates Notch1 signaling and influences cell regeneration, apoptosis and differentiation in carcinogenesis. ADAM17 by regulating the release and density of various leukocyte cell surface proteins modulates inflammation in acute pulmonary inflammation, lung injury and pulmonary fibrosis. Induced ADAM17 -mediated shedding is associated with cardiac and renal damage and pathogenesis of diabetes, atherosclerosis and heart failure. Interestingly, ADAM17 may mediate Shedding of the ACE2 in SARS infection, however only cleavage by TMPRSS2 results in augmented SARS-S-driven entry.**Ref:** [42–48];  **Gene Ontology:** |
| GP2 | 7 | 1.83 | 2.36 | 1.75 | **-** | **-** | **-** | **-** | **-** | **-** | **-** | **Description:** Pancreatic secretory granule membrane major glycoprotein GP2; Glycoprotein 2;  **Gene Ontology:** |
| LNPEP | 7 | 2.44 | 1.60 | 4.51 | **-** | **-** | **-** | **-** | **-** | **-** | **-** | **Description:** Insulin-regulated membrane aminopeptidase; Release of an N-terminal amino acid, cleaves before cysteine, leucine as well as other amino acids. Degrades peptide hormones such as oxytocin, vasopressin and angiotensin III, and plays a role in maintaining homeostasis during pregnancy. May be involved in the inactivation of neuronal peptides in the brain. Cleaves Met-enkephalin and dynorphin. Binds angiotensin IV and may be the angiotensin IV receptor in the brain; Aminopeptidases;  **Gene Ontology:** |
| AGTRAP | 6 | 2.22 | 2.36 | 4.47 | **-** | **-** | **-** | **-** | **-** | **-** | **-** | **Description:** Type-1 angiotensin II receptor-associated protein; Appears to be a negative regulator of type-1 angiotensin II receptor-mediated signaling by regulating receptor internalisation as well as mechanism of receptor desensitization such as phosphorylation. Induces also a decrease in cell proliferation and angiotensin II-stimulated transcriptional activity.  **Gene Ontology:** |
| FABP2 | 6 | 1.70 | 1.53 | 1.93 | **Y** | **-** | **Y** | **Y** | **Y** | **-** | **Y** | **Description:** Intestinal-type fatty acid-binding protein; FABP are thought to play a role in the intracellular transport of long-chain fatty acids and their acyl-CoA esters. FABP2 is probably involved in triglyceride-rich lipoprotein synthesis. Binds saturated long-chain fatty acids with a high affinity, but binds with a lower affinity to unsaturated long- chain fatty acids. FABP2 may also help maintain energy homeostasis by functioning as a lipid sensor; Belongs to the calycin superfamily. Fatty-acid binding protein (FABP) family.  **COVID-19 related function:** The protein encoded by this gene is an intracellular fatty acid-binding protein that participates in the uptake, intracellular metabolism, and transport of long-chain fatty acids. FABP2 may act as a lipid sensor to maintain energy homeostasis. It is involved in pathogenesis of insulin resistance, obesity and metabolic syndrome. Elevated serum concentrations of I-FABP were found in patients suffering from abdominal sepsis and systemic inflammatory response. Circulating I-FABP levels were associated with adverse clinical outcomes in patients with acute decompensated HF with systolic dysfunction and among patients with chronic renal failure. Polymorphisms of FABP2 in some populations were associated with incidence of atherosclerosis and T2DM and also with myocardial infarction among individuals with or without other risk factors including hypertension, hypercholesterolemia, or diabetes mellitus.**Ref:** [49–57];  **Gene Ontology:** |
| CALM1 | 6 | 4.02 | 4.86 | 5.00 | **-** | **-** | **-** | **-** | **-** | **-** | **-** | **Description:** Calmodulin 1 (phosphorylase kinase, delta); Calmodulin mediates the control of a large number of enzymes, ion channels, aquaporins and other proteins through calcium-binding. Among the enzymes to be stimulated by the calmodulin-calcium complex are a number of protein kinases and phosphatases. Together with CCP110 and centrin, is involved in a genetic pathway that regulates the centrosome cycle and progression through cytokinesis. Mediates calcium-dependent inactivation of CACNA1C. Positively regulates calcium-activated potassium channel activity of KCNN2.  **Gene Ontology:** cardiac muscle functions |
| CDK4 | 6 | 2.98 | 4.03 | 4.64 | **Y** | **-** | **-** | **-** | **-** | **-** | **-** | **Description:** Cell division protein kinase 4; Ser/Thr-kinase component of cyclin D-CDK4 (DC) complexes that phosphorylate and inhibit members of the retinoblastoma (RB) protein family including RB1 and regulate the cell-cycle during G(1)/S transition. Phosphorylation of RB1 allows dissociation of the transcription factor E2F from the RB/E2F complexes and the subsequent transcription of E2F target genes which are responsible for the progression through the G(1) phase. Hypophosphorylates RB1 in early G(1) phase. Cyclin D-CDK4 complexes are major integrators of various mitogenenic and antimitogenic signals. Also phosphorylates SMAD3 in a cell-cycle-dependent manner and represses its transcriptional activity. Component of the ternary complex, cyclin D/CDK4/CDKN1B, required for nuclear translocation and activity of the cyclin D-CDK4 complex.  **Gene Ontology:** fibrosis |
| CALM3 | 6 | 3.19 | 4.85 | 5.00 | **-** | **-** | **-** | **-** | **-** | **-** | **-** | **Description:** Calmodulin 3 (phosphorylase kinase, delta); Calmodulin mediates the control of a large number of enzymes, ion channels, aquaporins and other proteins through calcium-binding. Among the enzymes to be stimulated by the calmodulin-calcium complex are a number of protein kinases and phosphatases. Together with CCP110 and centrin, is involved in a genetic pathway that regulates the centrosome cycle and progression through cytokinesis. Mediates calcium-dependent inactivation of CACNA1C. Positively regulates calcium-activated potassium channel activity of KCNN2.  **Gene Ontology:** cardiac muscle functions |
| NPC1 | 6 | 2.00 | 2.16 | 4.90 | **-** | **-** | **-** | **-** | **-** | **-** | **Y** | **Description:** Niemann-Pick disease, type C1; Intracellular cholesterol transporter which acts in concert with NPC2 and plays an important role in the egress of cholesterol from the endosomal/lysosomal compartment. Both NPC1 and NPC2 function as the cellular 'tag team duo' (TTD) to catalyze the mobilization of cholesterol within the multivesicular environment of the late endosome (LE) to effect egress through the limiting bilayer of the LE. NPC2 binds unesterified cholesterol that has been released from LDLs in the lumen of the late endosomes/lysosomes and transfers it to the cholesterol-binding pocket of the N-terminal domain of NPC1. Cholesterol binds to NPC1 with the hydroxyl group buried in the binding pocket and is exported from the limiting membrane of late endosomes/ lysosomes to the ER and plasma membrane by an unknown mechanism. Binds oxysterol with higher affinity than cholesterol. May play a role in vesicular trafficking in glia, a process that may be crucial for maintaining the structural and functional integrity of nerve terminals; Belongs to the patched family.  **COVID-19 related function:** NPC protein mediates intracellular cholesterol trafficking and acts as an endosomal entry receptor for ebolavirus. Gene ontology enrichment analysis revealed that differentially expressed proteins in NPC1 knockout fibroblasts were associated with exacerbated inflammation, and oxidative stress. In experimental models loss of function of NCP1 results in dysregulated expression of genes associated with ROS formation and fibrosis, leading to organ damage. In line with this findings, oxidative stress pathway was also activated in sera of patients with mutation of NPC1 gene.**Ref:** [58–62];  **Gene Ontology:** |
| ANG | 6 | 2.88 | 3.01 | 2.86 | **-** | **-** | **-** | **-** | **Y** | **-** | **-** | **Description:** Angiogenin, ribonuclease, RNase A family, 5; Binds to actin on the surface of endothelial cells; once bound, angiogenin is endocytosed and translocated to the nucleus. Stimulates ribosomal RNA synthesis including that containing the initiation site sequences of 45S rRNA. Cleaves tRNA within anticodon loops to produce tRNA-derived stress-induced fragments (tiRNAs) which inhibit protein synthesis and triggers the assembly of stress granules (SGs). Angiogenin induces vascularization of normal and malignant tissues. Angiogenic activity is regulated by interaction with RNH1 in vivo; Ribonuclease A family;  **Gene Ontology:** angiogenesis |
| LTA4H | 6 | 3.06 | 4.85 | 4.76 | **-** | **-** | **-** | **-** | **-** | **-** | **-** | **Description:** Leukotriene A(4) hydrolase; Epoxide hydrolase that catalyzes the final step in the biosynthesis of the proinflammatory mediator leukotriene B4. Has also aminopeptidase activity; M1 metallopeptidases;  **Gene Ontology:** |
| GHRL | 6 | 4.53 | 1.80 | 4.20 | **-** | **Y** | **Y** | **-** | **Y** | **-** | **-** | **Description:** Growth hormone-releasing peptide; Ghrelin is the ligand for growth hormone secretagogue receptor type 1 (GHSR). Induces the release of growth hormone from the pituitary. Has an appetite-stimulating effect, induces adiposity and stimulates gastric acid secretion. Involved in growth regulation; Endogenous ligands;  **Gene Ontology:** angiogenesis; inflammation |
| LCT | 6 | 2.21 | 2.27 | 2.38 | **-** | **-** | **-** | **-** | **-** | **-** | **-** | **Description:** Lactase-phlorizin hydrolase; LPH splits lactose in the small intestine.  **Gene Ontology:** |
| CALM2 | 5 | 3.63 | 4.95 | 5.00 | **-** | **-** | **-** | **-** | **-** | **-** | **-** | **Description:** Calmodulin 2 (phosphorylase kinase, delta); EF-hand domain containing;  **Gene Ontology:** cardiac muscle functions |
| VIL1 | 5 | 1.94 | 2.08 | 2.54 | **-** | **-** | **-** | **-** | **-** | **Y** | **-** | **Description:** Villin 1; Epithelial cell-specific Ca(2+)-regulated actin- modifying protein that modulates the reorganization of microvillar actin filaments. Plays a role in the actin nucleation, actin filament bundle assembly, actin filament capping and severing. Binds phosphatidylinositol 4,5-bisphosphate (PIP2) and lysophosphatidic acid (LPA); binds LPA with higher affinity than PIP2. Binding to LPA increases its phosphorylation by SRC and inhibits all actin-modifying activities. Binding to PIP2 inhibits actin-capping and -severing activities but enhances actin-bundling activity. Regulates the intestinal epithelial cell morphology, cell invasion, cell migration and apoptosis. Protects against apoptosis induced by dextran sodium sulfate (DSS) in the gastrointestinal epithelium. Appears to regulate cell death by maintaining mitochondrial integrity. Enhances hepatocyte growth factor (HGF)-induced epithelial cell motility, chemotaxis and wound repair. Upon S.flexneri cell infection, its actin-severing activity enhances actin-based motility of the bacteria and plays a role during the dissemination; Gelsolin/villins;  **Gene Ontology:** |
| APOA4 | 5 | 3.34 | 1.89 | 4.42 | **-** | **-** | **-** | **-** | **-** | **-** | **-** | **Description:** Apolipoprotein A-IV; May have a role in chylomicrons and VLDL secretion and catabolism. Required for efficient activation of lipoprotein lipase by ApoC-II; potent activator of LCAT. Apoa-IV is a major component of HDL and chylomicrons; Belongs to the apolipoprotein A1/A4/E family.  **Gene Ontology:** |
| KLK1 | 4 | 2.20 | 1.63 | 2.07 | **-** | **-** | **-** | **-** | **-** | **-** | **-** | **Description:** Kidney/pancreas/salivary gland kallikrein; Glandular kallikreins cleave Met-Lys and Arg-Ser bonds in kininogen to release Lys-bradykinin; Belongs to the peptidase S1 family. Kallikrein subfamily.  **Gene Ontology:** |
| CPA3 | 4 | 1.92 | 4.52 | 1.57 | **-** | **-** | **-** | **-** | **-** | **-** | **-** | **Description:** Carboxypeptidase A3 (mast cell); M14 carboxypeptidases;  **Gene Ontology:** |
| CTSA | 4 | 3.19 | 4.16 | 4.97 | **-** | **-** | **-** | **-** | **-** | **-** | **-** | **Description:** Protective protein for beta-galactosidase; Protective protein appears to be essential for both the activity of beta-galactosidase and neuraminidase, it associates with these enzymes and exerts a protective function necessary for their stability and activity. This protein is also a carboxypeptidase and can deamidate tachykinins.  **Gene Ontology:** |
| TREH | 4 | 0.81 | 0.86 | 1.69 | **-** | **-** | **-** | **-** | **-** | **-** | **-** | **Description:** Trehalase (brush-border membrane glycoprotein); Intestinal trehalase is probably involved in the hydrolysis of ingested trehalose; Belongs to the glycosyl hydrolase 37 family.  **Gene Ontology:** |
| SLC6A19 | 4 | 1.01 | 0.66 | 1.40 | **-** | **-** | **-** | **-** | **-** | **-** | **-** | **Description:** Solute carrier family 6 (neutral amino acid transporter), member 19; Transporter that mediates resorption of neutral amino acids across the apical membrane of renal and intestinal epithelial cells. This uptake is sodium-dependent and chloride- independent; Belongs to the sodium:neurotransmitter symporter (SNF) (TC 2.A.22) family. SLC6A19 subfamily.  **Gene Ontology:** |
| SLC15A2 | 3 | 1.00 | 2.17 | 4.90 | **-** | **-** | **-** | **-** | **-** | **-** | **-** | **Description:** Solute carrier family 15 (oligopeptide transporter), member 2; Proton-coupled intake of oligopeptides of 2 to 4 amino acids with a preference for dipeptides; Belongs to the PTR2/POT transporter (TC 2.A.17) family.  **Gene Ontology:** |
| CLEC4M | 3 | 2.07 | 1.99 | 4.32 | **-** | **-** | **-** | **-** | **-** | **-** | **Y** | **Description:** Dendritic cell-specific ICAM-3-grabbing non-integrin 2; Probable pathogen-recognition receptor involved in peripheral immune surveillance in liver. May mediate the endocytosis of pathogens which are subsequently degraded in lysosomal compartments. Is a receptor for ICAM3, probably by binding to mannose-like carbohydrates; CD molecules;  **COVID-19 related function:** C-type lectin, also known as CD209L, L-Sign functions in cell adhesion and pathogen recognition. It is involved in pathogenesis of several viral infections including Ebola, Hepatitis C-Virus, Hiv-1, CoV-229E, HHV-5 and influenzavirus. CD209L might mediate transfer of HIV1 virus to dendritic or T cells. Moreover, this protein is recognized as a receptor for severe acute respiratory syndrome coronavirus. CD209L, perhaps in combination with ACE2, in lymph nodes, Peyer's patches, and lung may contribute to the spread of SARS-COV infection.**Ref:** [63–65];  **Gene Ontology:** |
| NPC1L1 | 3 | 1.48 | 0.53 | 1.57 | **-** | **-** | **-** | **-** | **-** | **-** | **-** | **Description:** Niemann-Pick C1-like protein 1; Plays a major role in cholesterol homeostasis. Is critical for the uptake of cholesterol across the plasma membrane of the intestinal enterocyte. Is the direct molecular target of ezetimibe, a drug that inhibits cholesterol absorption. Lack of activity leads to multiple lipid transport defects. The protein may have a function in the transport of multiple lipids and their homeostasis, and may play a critical role in regulating lipid metabolism. Acts as a negative regulator of NPC2 and down- regulates its expression and secretion by inhibiting its maturation and accelerating its degradation; Solute carriers;  **Gene Ontology:** |
| SOX14 | 2 |  |  | 4.32 | **-** | **-** | **-** | **-** | **-** | **-** | **-** | **Description:** SRY (sex determining region Y)-box 14; Acts as a negative regulator of transcription; SRY-boxes;  **Gene Ontology:** |
| KLK2 | 2 | 1.31 | 1.15 | 2.38 | **-** | **-** | **-** | **-** | **-** | **-** | **-** | **Description:** Kallikrein-related peptidase 2; Glandular kallikreins cleave Met-Lys and Arg-Ser bonds in kininogen to release Lys-bradykinin.  **Gene Ontology:** |
| MRGPRD | 2 | 2.00 | 1.02 | 2.65 | **-** | **-** | **-** | **-** | **-** | **-** | **-** | **Description:** Mas-related G-protein coupled receptor member D; May regulate nociceptor function and/or development, including the sensation or modulation of pain. Functions as a specific membrane receptor for beta-alanine. Beta-alanine at micromolar doses specifically evoked Ca(2+) influx in cells expressing the receptor. Beta-alanine decreases forskolin- stimulated cAMP production in cells expressing the receptor, suggesting that the receptor couples with G-protein G(q) and G(i); G protein-coupled receptors, Class A orphans;  **Gene Ontology:** |
| TMPRSS2 | 2 | 1.54 | 2.58 | 4.31 | **-** | **-** | **-** | **-** | **-** | **-** | **Y** | **Description:** Transmembrane protease, serine 2; Serine protease that proteolytically cleaves and activates the viral spike glycoproteins which facilitate virus- cell membrane fusions; spike proteins are synthesized and maintained in precursor intermediate folding states and proteolysis permits the refolding and energy release required to create stable virus-cell linkages and membrane coalescence. Facilitates human SARS coronavirus (SARS-CoV) infection via two independent mechanisms, proteolytic cleavage of ACE2, which might promote viral uptake, and cleavage of coronavirus spike glycoprotein which activates the glycoprotein for cathepsin L- independent host cell entry. Proteolytically cleaves and activates the spike glycoproteins of human coronavirus 229E (HCoV-229E) and human coronavirus EMC (HCoV-EMC) and the fusion glycoproteins F0 of Sendai virus (SeV), human metapneumovirus (HMPV), human parainfluenza 1, 2, 3, 4a and 4b viruses (HPIV). Essential for spread and pathogenesis of influenza A virus (strains H1N1, H3N2 and H7N9); involved in proteolytic cleavage and activation of hemagglutinin (HA) protein which is essential for viral infectivity; Belongs to the peptidase S1 family.  **COVID-19 related function:** TMPRSS2 is a serine cellular protease known to be involved in spread of several viruses including influenza A viruses and coronaviruses. Similar to previous Sars-COV, Sars-CoV-2 uses ACE2 and TMPRSS2 for entry into target cells which is facilitated by the spike protein. TMPRSS2 cleaves and activates the coronavirus spike protein. Sars-CoV-2 spike protein priming by TMPRSS2 is essential for viral entry into target cells and for viral spread in the infected host, thus high TMPRSS2-expressing cells are potentially the most susceptible for Sars-Cov-2 infection.**Ref:** [66–70];  **Gene Ontology:** |
| TM4SF5 | 1 | 1.75 | 0.78 | 0.28 | **-** | **-** | **-** | **-** | **-** | **-** | **-** | **Description:** Transmembrane 4 L six family member 5; Belongs to the L6 tetraspanin family.  **Gene Ontology:** |
| SWI5 | 1 | 1.13 | 0.92 | 4.30 | **-** | **-** | **-** | **-** | **-** | **-** | **-** | **Description:** SWI5 recombination repair homolog (yeast); Component of the SWI5-SFR1 complex, a complex required for double-strand break repair via homologous recombination.  **Gene Ontology:** |
| SOX3 | 1 | 2.25 | 0.95 | 4.57 | **-** | **-** | **-** | **-** | **-** | **-** | **-** | **Description:** SRY (sex determining region Y)-box 3; Transcription factor required during the formation of the hypothalamo-pituitary axis. May function as a switch in neuronal development. Keeps neural cells undifferentiated by counteracting the activity of proneural proteins and suppresses neuronal differentiation. Required also within the pharyngeal epithelia for craniofacial morphogenesis. Controls a genetic switch in male development. Is necessary for initiating male sex determination by directing the development of supporting cell precursors (pre-Sertoli cells) as Sertoli rather than granulosa cells (By similarity); SRY-boxes;  **Gene Ontology:** |
| AAMP | 1 | 2.28 | 2.58 | 4.90 | **-** | **-** | **-** | **-** | **-** | **Y** | **-** | **Description:** Angio-associated, migratory cell protein; Plays a role in angiogenesis and cell migration. In smooth muscle cell migration, may act through the RhoA pathway; WD repeat domain containing;  **Gene Ontology:** angiogenesis |
| ENPP7 | 1 | 1.33 | 0.71 | 1.03 | **-** | **-** | **-** | **-** | **-** | **-** | **-** | **Description:** Ectonucleotide pyrophosphatase/phosphodiesterase family member 7; Converts sphingomyelin to ceramide. Also has phospholipase C activity toward palmitoyl lyso-phosphocholine. Does not appear to have nucleotide pyrophosphatase activity; Belongs to the nucleotide pyrophosphatase/phosphodiesterase family.  **Gene Ontology:** |
| CDHR2 | 0 | 0.97 | 0.67 | 0.86 | **-** | **-** | **-** | **-** | **-** | **-** | **-** | **Description:** Cadherin-related family member 2; Intermicrovillar adhesion molecule that forms, via its extracellular domain, calcium-dependent heterophilic complexes with CDHR5 on adjacent microvilli. Thereby, controls the packing of microvilli at the apical membrane of epithelial cells. Through its cytoplasmic domain, interacts with microvillus cytoplasmic proteins to form the intermicrovillar adhesion complex/IMAC. This complex plays a central role in microvilli and epithelial brush border differentiation. May also play a role in cell-cell adhesion and contact inhibition in epithelial cells; Cadherin related;  **Gene Ontology:** |
| MS4A10 | 0 |  | 0.55 | 0.55 | **-** | **-** | **-** | **-** | **-** | **-** | **-** | **Description:** Membrane-spanning 4-domains, subfamily A, member 10; May be involved in signal transduction as a component of a multimeric receptor complex; Membrane spanning 4-domains;  **Gene Ontology:** |

ACE2: Angiotensin-converting enzyme 2; TMPRSS2: Transmembrane protease, serine 2; RAAS: Renin-Angiotensin-Aldosterone System; NF-κB: nuclear factor kappa-light-chain-enhancer of activated B cells; MAPK: mitogen-activated protein kinases; MAS: MAS1 oncogene; Ang-(1-7): Angiotensin (1-7); AT_1:_ Angiotensin II receptor type 1; ERK1: extracellular signal-regulated kinase 1; TGF-β: Transforming growth factor beta; SMAD: Sma and Mad proteins from Caenorhabditis elegans and Drosophila; PKC: protein kinase C; PI3K-Akt: phosphatidylinositol 3-kinase: eNOS: Endothelial NOS; JNK1/2: c-Jun N-terminal kinase ½; MMP: matrix metalloproteinase; NADPH oxidase: nicotinamide adenine dinucleotide phosphate oxidase; SARS: Severe Acute Respiratgory Syndrome; CoV: coronavirus; ROS: reactive oxygen species; T2DM: Type 2 diabetes mellitus; COVID-19: Coronavirus disease 2019;

**REFERENCES**

1.	Montezano AC, Cat AND, Rios FJ, Touyz RM. Angiotensin II and Vascular Injury. Current Hypertension Reports. 2014.

2.	Muus C, Luecken MD, Eraslan G, Waghray A, Heimberg G, Sikkema L, Kobayashi Y, Vaishnav ED, Subramanian A, Smilie C, Jagadeesh K, Duong ET, Fiskin E, Triglia ET, Ansari M, Cai P, Lin B, Buchanan J, Chen S, Shu J, Haber AL, Chung H, Montoro DT, Adams T, Aliee H, Samuel J, Andrusivova AZ, Angelidis I, Ashenberg O, Bassler K, et al. Integrated analyses of single-cell atlases reveal age, gender, and smoking status associations with cell type-specific expression of mediators of SARS-CoV-2 viral entry and highlights inflammatory programs in putative target cells.

3.	Crackower MA, Sarao R, Oudit GY, Yagil C, Kozieradzki I, Scanga SE, Oliveira-dos-Santos AJ, Costa J da, Zhang L, Pei Y, Scholey J, Ferrario CM, Manoukian AS, Chappell MC, Backx PH, Yagil Y, Penninger JM. Angiotensin-converting enzyme 2 is an essential regulator of heart function. *Nature* 2002;**417**:822–828.

4.	Xue T, Wei N, Xin Z, Qingyu X. Angiotensin-converting enzyme-2 overexpression attenuates inflammation in rat model of chronic obstructive pulmonary disease. *Inhal Toxicol* 2014;**26**:14–22.

5.	Silva ACS e., Silveira KD, Ferreira AJ, Teixeira MM. ACE2, angiotensin-(1-7) and Mas receptor axis in inflammation and fibrosis. British Journal of Pharmacology. 2013. p. 477–492.

6.	Patel VB, Mori J, McLean BA, Basu R, Das SK, Ramprasath T, Parajuli N, Penninger JM, Grant MB, Lopaschuk GD, Oudit GY. ACE2 Deficiency Worsens Epicardial Adipose Tissue Inflammation and Cardiac Dysfunction in Response to Diet-Induced Obesity. *Diabetes* 2016;**65**:85–95.

7.	Liu Z, Huang XR, Chen H-Y, Penninger JM, Lan HY. Loss of angiotensin-converting enzyme 2 enhances TGF-β/Smad-mediated renal fibrosis and NF-κB-driven renal inflammation in a mouse model of obstructive nephropathy. Laboratory Investigation. 2012. p. 650–661.

8.	Kassiri Z, Zhong J, Guo D, Basu R, Wang X, Liu PP, Scholey JW, Penninger JM, Oudit GY. Loss of angiotensin-converting enzyme 2 accelerates maladaptive left ventricular remodeling in response to myocardial infarction. *Circ Heart Fail* 2009;**2**:446–455.

9.	Patel VB, Zhong J-C, Grant MB, Oudit GY. Role of the ACE2/Angiotensin 1–7 Axis of the Renin–Angiotensin System in Heart Failure. Circulation Research. 2016. p. 1313–1326.

10.	Hamming I, Cooper ME, Haagmans BL, Hooper NM, Korstanje R, Osterhaus A, Timens W, Turner AJ, Navis G, Goor H van. The emerging role of ACE2 in physiology and disease. The Journal of Pathology. 2007. p. 1–11.

11.	Basu R, Poglitsch M, Yogasundaram H, Thomas J, Rowe BH, Oudit GY. Roles of Angiotensin Peptides and Recombinant Human ACE2 in Heart Failure. *J Am Coll Cardiol* 2017;**69**:805–819.

12.	Zhong J, Basu R, Guo D, Chow FL, Byrns S, Schuster M, Loibner H, Wang X-H, Penninger JM, Kassiri Z, Oudit GY. Angiotensin-converting enzyme 2 suppresses pathological hypertrophy, myocardial fibrosis, and cardiac dysfunction. *Circulation* 2010;**122**:717–728, 18 p following 728.

13.	Imai Y, Kuba K, Rao S, Huan Y, Guo F, Guan B, Yang P, Sarao R, Wada T, Leong-Poi H, Crackower MA, Fukamizu A, Hui C-C, Hein L, Uhlig S, Slutsky AS, Jiang C, Penninger JM. Angiotensin-converting enzyme 2 protects from severe acute lung failure. Nature. 2005. p. 112–116.

14.	Röhrborn D. DPP4 in diabetes. Frontiers in Immunology. 2015.

15.	Ervinna N, Mita T, Yasunari E, Azuma K, Tanaka R, Fujimura S, Sukmawati D, Nomiyama T, Kanazawa A, Kawamori R, Fujitani Y, Watada H. Anagliptin, a DPP-4 inhibitor, suppresses proliferation of vascular smooth muscles and monocyte inflammatory reaction and attenuates atherosclerosis in male apo E-deficient mice. *Endocrinology* 2013;**154**:1260–1270.

16.	Fadini GP, Avogaro A. Cardiovascular effects of DPP-4 inhibition: beyond GLP-1. *Vascul Pharmacol* 2011;**55**:10–16.

17.	Wang N, Shi X, Jiang L, Zhang S, Wang D, Tong P, Guo D, Fu L, Cui Y, Liu X, Arledge KC, Chen Y-H, Zhang L, Wang X. Structure of MERS-CoV spike receptor-binding domain complexed with human receptor DPP4. *Cell Res* 2013;**23**:986–993.

18.	Steinbrecher A, Reinhold D, Quigley L, Gado A, Tresser N, Izikson L, Born I, Faust J, Neubert K, Martin R, Ansorge S, Brocke S. Targeting Dipeptidyl Peptidase IV (CD26) Suppresses Autoimmune Encephalomyelitis and Up-Regulates TGF-β1 Secretion In Vivo. The Journal of Immunology. 2001. p. 2041–2048.

19.	Wronkowitz N, Görgens SW, Romacho T, Villalobos LA, Sánchez-Ferrer CF, Peiró C, Sell H, Eckel J. Soluble DPP4 induces inflammation and proliferation of human smooth muscle cells via protease-activated receptor 2. Biochimica et Biophysica Acta (BBA) - Molecular Basis of Disease. 2014. p. 1613–1621.

20.	Barchetta I, Cavallo MG, Baroni MG. COVID-19 and diabetes: Is this association driven by the DPP4 receptor? Potential clinical and therapeutic implications. Diabetes Research and Clinical Practice. 2020. p. 108165.

21.	Kawasaki T, Chen W, Htwe YM, Tatsumi K, Dudek SM. DPP4 inhibition by sitagliptin attenuates LPS-induced lung injury in mice. American Journal of Physiology-Lung Cellular and Molecular Physiology. 2018. p. L834–L845.

22.	Lietzau G, Davidsson W, Östenson C-G, Chiazza F, Nathanson D, Pintana H, Skogsberg J, Klein T, Nyström T, Darsalia V, Patrone C. Type 2 diabetes impairs odour detection, olfactory memory and olfactory neuroplasticity; effects partly reversed by the DPP-4 inhibitor Linagliptin. Acta Neuropathologica Communications. 2018.

23.	Reinhold D, Biton A, Pieper S, Lendeckel U, Faust J, Neubert K, Bank U, Täger M, Ansorge S, Brocke S. Dipeptidyl peptidase IV (DP IV, CD26) and aminopeptidase N (APN, CD13) as regulators of T cell function and targets of immunotherapy in CNS inflammation. International Immunopharmacology. 2006. p. 1935–1942.

24.	Pérez I, Varona A, Blanco L, Gil J, Santaolalla F, Zabala A, Ibarguen AM, Irazusta J, Larrinaga G. Increased APN/CD13 and acid aminopeptidase activities in head and neck squamous cell carcinoma. Head & Neck. 2009. p. 1335–1340.

25.	Mina-Osorio P. The moonlighting enzyme CD13: old and new functions to target. Trends in Molecular Medicine. 2008. p. 361–371.

26.	Fukasawa K, Fujii H, Saitoh Y, Koizumi K, Aozuka Y, Sekine K, Yamada M, Saiki I, Nishikawa K. Aminopeptidase N (APN/CD13) is selectively expressed in vascular endothelial cells and plays multiple roles in angiogenesis. Cancer Letters. 2006. p. 135–143.

27.	Lu C, Amin MA, Fox DA. CD13/Aminopeptidase N Is a Potential Therapeutic Target for Inflammatory Disorders. The Journal of Immunology. 2020. p. 3–11.

28.	Danziger RS. Aminopeptidase N in arterial hypertension. *Heart Fail Rev* 2008;**13**:293–298.

29.	Deshmane SL, Kremlev S, Amini S, Sawaya BE. Monocyte Chemoattractant Protein-1 (MCP-1): An Overview. Journal of Interferon & Cytokine Research. 2009. p. 313–326.

30.	Cheung CY, Poon LLM, Ng IHY, Luk W, Sia S-F, Wu MHS, Chan K-H, Yuen K-Y, Gordon S, Guan Y, Peiris JSM. Cytokine Responses in Severe Acute Respiratory Syndrome Coronavirus-Infected Macrophages In Vitro: Possible Relevance to Pathogenesis. Journal of Virology. 2005. p. 7819–7826.

31.	Wong CK, Lam CWK, Wu AKL, Ip WK, Lee NLS, Chan IHS, Lit LCW, Hui DSC, Chan MHM, Chung SSC, Sung JJY. Plasma inflammatory cytokines and chemokines in severe acute respiratory syndrome. Clinical & Experimental Immunology. 2004. p. 95–103.

32.	Huang K-J, Su I-J, Theron M, Wu Y-C, Lai S-K, Liu C-C, Lei H-Y. An interferon-?-related cytokine storm in SARS patients. Journal of Medical Virology. 2005. p. 185–194.

33.	Lin J, Kakkar V, Lu X. Impact of MCP -1 in Atherosclerosis. Current Pharmaceutical Design. 2014. p. 4580–4588.

34.	Cameron MJ, Bermejo-Martin JF, Danesh A, Muller MP, Kelvin DJ. Human immunopathogenesis of severe acute respiratory syndrome (SARS). *Virus Res* 2008;**133**:13–19.

35.	Chen I-Y, Chang SC, Wu H-Y, Yu T-C, Wei W-C, Lin S, Chien C-L, Chang M-F. Upregulation of the Chemokine (C-C Motif) Ligand 2 via a Severe Acute Respiratory Syndrome Coronavirus Spike-ACE2 Signaling Pathway. Journal of Virology. 2010. p. 7703–7712.

36.	Rose CE, Edward Rose C, Sung S-SJ, Fu SM. Significant Involvement of CCL2 (MCP-1) in Inflammatory Disorders of the Lung. Microcirculation. 2003. p. 273–288.

37.	Jabara HH, Boyden SE, Chou J, Ramesh N, Massaad MJ, Benson H, Bainter W, Fraulino D, Rahimov F, Sieff C, Liu Z-J, Alshemmari SH, Al-Ramadi BK, Al-Dhekri H, Arnaout R, Abu-Shukair M, Vatsayan A, Silver E, Ahuja S, Davies EG, Sola-Visner M, Ohsumi TK, Andrews NC, Notarangelo LD, Fleming MD, Al-Herz W, Kunkel LM, Geha RS. A missense mutation in TFRC, encoding transferrin receptor 1, causes combined immunodeficiency. *Nat Genet* 2016;**48**:74–78.

38.	Broder C, Becker-Pauly C. The metalloproteases meprin α and meprin β: unique enzymes in inflammation, neurodegeneration, cancer and fibrosis. *Biochem J* 2013;**450**:253–264.

39.	Lam UDP, Lerchbaum E, Schweighofer N, Trummer O, Eberhard K, Genser B, Pieber TR, Obermayer-Pietsch B. Association of MEP1A gene variants with insulin metabolism in central European women with polycystic ovary syndrome. Gene. 2014. p. 245–252.

40.	Mathew R, Futterweit S, Valderrama E, Tarectecan AA, Bylander JE, Bond JS, Trachtman H. Meprin-α in chronic diabetic nephropathy: interaction with the renin-angiotensin axis. American Journal of Physiology-Renal Physiology. 2005. p. F911–F921.

41.	Prox J, Arnold P, Becker-Pauly C. Meprin α and meprin β: Procollagen proteinases in health and disease. Matrix Biology. 2015. p. 7–13.

42.	Arndt PG, Strahan B, Wang Y, Long C, Horiuchi K, Walcheck B. Leukocyte ADAM17 regulates acute pulmonary inflammation. *PLoS One* 2011;**6**:e19938.

43.	Scheller J, Chalaris A, Garbers C, Rose-John S. ADAM17: a molecular switch to control inflammation and tissue regeneration. *Trends Immunol* 2011;**32**:380–387.

44.	Chen J-Y, Lin C-H, Chen B-C. Hypoxia-induced ADAM 17 expression is mediated by RSK1-dependent C/EBPβ activation in human lung fibroblasts. Molecular Immunology. 2017. p. 155–163.

45.	Kefaloyianni E, Muthu ML, Kaeppler J, Sun X, Sabbisetti V, Chalaris A, Rose-John S, Wong E, Sagi I, Waikar SS, Rennke H, Humphreys BD, Bonventre JV, Herrlich A. ADAM17 substrate release in proximal tubule drives kidney fibrosis. JCI Insight. 2016.

46.	Menghini R, Fiorentino L, Casagrande V, Lauro R, Federici M. The role of ADAM17 in metabolic inflammation. *Atherosclerosis* 2013;**228**:12–17.

47.	Arribas J, Esselens C. ADAM17 as a therapeutic target in multiple diseases. *Curr Pharm Des* 2009;**15**:2319–2335.

48.	Heurich A, Hofmann-Winkler H, Gierer S, Liepold T, Jahn O, Pohlmann S. TMPRSS2 and ADAM17 Cleave ACE2 Differentially and Only Proteolysis by TMPRSS2 Augments Entry Driven by the Severe Acute Respiratory Syndrome Coronavirus Spike Protein. Journal of Virology. 2014. p. 1293–1307.

49.	Atshaves BP, Foxworth WB, Frolov A, Roths JB, Kier AB, Oetama BK, Piedrahita JA, Schroeder F. Cellular differentiation and I-FABP protein expression modulate fatty acid uptake and diffusion. American Journal of Physiology-Cell Physiology. 1998. p. C633–C644.

50.	Veerkamp JH, Peeters RA, R G H. Structural and functional features of different types of cytoplasmic fatty acid-binding proteins. Biochimica et Biophysica Acta (BBA) - Lipids and Lipid Metabolism. 1991. p. 1–24.

51.	Albala C, Santos JL, Cifuentes M, Villarroel AC, Lera L, Liberman C, Angel B, Pérez-Bravo F. Intestinal FABP2 A54T polymorphism: association with insulin resistance and obesity in women. *Obes Res* 2004;**12**:340–345.

52.	Vimaleswaran KS, Radha V, Mohan V. Thr54 allele carriers of the Ala54Thr variant of FABP2 gene have associations with metabolic syndrome and hypertriglyceridemia in urban South Indians. *Metabolism* 2006;**55**:1222–1226.

53.	Kitai T, Kim Y-H, Kiefer K, Morales R, Borowski AG, Grodin JL, Wilson Tang WH. Circulating intestinal fatty acid-binding protein (I-FABP) levels in acute decompensated heart failure. Clinical Biochemistry. 2017. p. 491–495.

54.	Okada K, Sekino M, Funaoka H, Sato S, Ichinomiya T, Murata H, Maekawa T, Nishikido M, Eishi K, Hara T. Intestinal fatty acid–binding protein levels in patients with chronic renal failure. Journal of Surgical Research. 2018. p. 94–100.

55.	Khattab SA, Abo-Elmatty DM, Ghattas MH, Mesbah NM, Mehanna ET. Intestinal fatty acid binding protein Ala54Thr polymorphism is associated with peripheral atherosclerosis combined with type 2 diabetes mellitus. Journal of Diabetes. 2017. p. 821–826.

56.	Qiu C-J, Ye X-Z, Yu X-J, Peng X-R, Li T-H. Association between FABP2 Ala54Thr polymorphisms and type 2 diabetes mellitus risk: a HuGE Review and Meta-Analysis. Journal of Cellular and Molecular Medicine. 2014. p. 2530–2535.

57.	Nishihama K, Yamada Y, Matsuo H, Segawa T, Watanabe S, Kato K, Yajima K, Hibino T, Yokoi K, Ichihara S, Metoki N, Yoshida H, Satoh K, Nozawa Y. Association of gene polymorphisms with myocardial infarction in individuals with or without conventional coronary risk factors. International Journal of Molecular Medicine. 2007.

58.	Kwon HJ, Abi-Mosleh L, Wang ML, Deisenhofer J, Goldstein JL, Brown MS, Infante RE. Structure of N-Terminal Domain of NPC1 Reveals Distinct Subdomains for Binding and Transfer of Cholesterol. Cell. 2009. p. 1213–1224.

59.	Scott C, Ioannou YA. The NPC1 protein: structure implies function. Biochimica et Biophysica Acta (BBA) - Molecular and Cell Biology of Lipids. 2004. p. 8–13.

60.	Rauniyar N, Subramanian K, Lavallée-Adam M, Martínez-Bartolomé S, Balch WE, Yates JR 3rd. Quantitative Proteomics of Human Fibroblasts with I1061T Mutation in Niemann-Pick C1 (NPC1) Protein Provides Insights into the Disease Pathogenesis. *Mol Cell Proteomics* 2015;**14**:1734–1749.

61.	Cluzeau CVM, Watkins-Chow DE, Fu R, Borate B, Yanjanin N, Dail MK, Davidson CD, Walkley SU, Ory DS, Wassif CA, Pavan WJ, Porter FD. Microarray expression analysis and identification of serum biomarkers for Niemann–Pick disease, type C1. Human Molecular Genetics. 2012. p. 3632–3646.

62.	Fu R, Yanjanin NM, Bianconi S, Pavan WJ, Porter FD. Oxidative stress in Niemann–Pick disease, type C. Molecular Genetics and Metabolism. 2010. p. 214–218.

63.	Jeffers SA, Tusell SM, Gillim-Ross L, Hemmila EM, Achenbach JE, Babcock GJ, Thomas WD, Thackray LB, Young MD, Mason RJ, Ambrosino DM, Wentworth DE, DeMartini JC, Holmes KV. CD209L (L-SIGN) is a receptor for severe acute respiratory syndrome coronavirus. Proceedings of the National Academy of Sciences. 2004. p. 15748–15753.

64.	Gillespie L, Roosendahl P, Ng WC, Brooks AG, Reading PC, Londrigan SL. Endocytic function is critical for influenza A virus infection via DC-SIGN and L-SIGN. Scientific Reports. 2016.

65.	Jameson B, Baribaud F, Pöhlmann S, Ghavimi D, Mortari F, Doms RW, Iwasaki A. Expression of DC-SIGN by dendritic cells of intestinal and genital mucosae in humans and rhesus macaques. *J Virol* 2002;**76**:1866–1875.

66.	Kawase M, Shirato K, Hoek L van der, Taguchi F, Matsuyama S. Simultaneous treatment of human bronchial epithelial cells with serine and cysteine protease inhibitors prevents severe acute respiratory syndrome coronavirus entry. *J Virol* 2012;**86**:6537–6545.

67.	Hoffmann M, Kleine-Weber H, Krüger N, Müller M, Drosten C, Pöhlmann S. The novel coronavirus 2019 (2019-nCoV) uses the SARS-coronavirus receptor ACE2 and the cellular protease TMPRSS2 for entry into target cells.

68.	Hoffmann M, Kleine-Weber H, Schroeder S, Krüger N, Herrler T, Erichsen S, Schiergens TS, Herrler G, Wu N-H, Nitsche A, Müller MA, Drosten C, Pöhlmann S. SARS-CoV-2 Cell Entry Depends on ACE2 and TMPRSS2 and Is Blocked by a Clinically Proven Protease Inhibitor. Cell. 2020. p. 271–280.e8.

69.	Matsuyama S, Nao N, Shirato K, Kawase M, Saito S, Takayama I, Nagata N, Sekizuka T, Katoh H, Kato F, Sakata M, Tahara M, Kutsuna S, Ohmagari N, Kuroda M, Suzuki T, Kageyama T, Takeda M. Enhanced isolation of SARS-CoV-2 by TMPRSS2-expressing cells. Proceedings of the National Academy of Sciences. 2020. p. 7001–7003.

70.	Iwata-Yoshikawa N, Okamura T, Shimizu Y, Hasegawa H, Takeda M, Nagata N. TMPRSS2 Contributes to Virus Spread and Immunopathology in the Airways of Murine Models after Coronavirus Infection. *J Virol* 2019;**93**.

71.	Aroor AR, Mandavia CH, Sowers JR. Insulin Resistance and Heart Failure. Heart Failure Clinics. 2012. p. 609–617.

72.	Bertrand L, Horman S, Beauloye C, Vanoverschelde J-L. Insulin signalling in the heart. Cardiovascular Research. 2008. p. 238–248.

73.	Montagnani M, Quon MJ. Insulin action in vascular endothelium: potential mechanisms linking insulin resistance with hypertension. Diabetes, Obesity and Metabolism. 2000. p. 285–292.

74.	Adler A. Obesity and target organ damage: diabetes. International Journal of Obesity. 2002. p. S11–S14.

75.	Folli F, Saad MJA, Velloso L, Hansen H, Carandente O, Feener E, Kahn C. Crosstalk between insulin and angiotensin II signalling systems. Experimental and Clinical Endocrinology & Diabetes. 2009. p. 133–139.
